# Bayesian inference of functional asymmetry in a ligand-gated ion channel

**DOI:** 10.1101/2025.06.20.660684

**Authors:** Luciano Moffatt, Gustavo Pierdominici-Sottile

## Abstract

Ligand-gated ion channels enable rapid cellular signaling by coupling extracellular cues to conformational transitions and ionic fluxes[1–5]. ATP-activated P2X receptors, with their minimalist trimeric architecture, serve as models for studying allosteric activation[6–8]. Although high-resolution structures reveal closed and open states[8, 9], the physical basis of the “flip state” and the emergence of negative cooperativity remain unresolved. Here we combine a recursive Bayesian framework (MacroIR) applied to outside-out patch-clamp recordings, with molecular dynamics simulations, to show that P2×2 activation proceeds via a directional, asymmetric coupling mechanism. ATP binding selectively lowers the energetic barrier for rotation of one subunit at the binding interface, promoting partial activation and substantial conductance, while minimally affecting its neighbor. This rotation, in turn, increases the barrier for subsequent ATP binding, providing a mechanistic explanation for negative cooperativity[7]. In the absence of ligand, elevated barriers stabilize the closed state, quantitatively accounting for spontaneous current fluctuations. These findings overturn the prevailing assumption of symmetric, concerted activation, and demonstrate that the classical flip state arises as a necessary physical intermediate. By showing that ligand-induced modulation of activation barriers can drive symmetry breaking in homomeric channels, our results establish a general principle for dynamic protein assemblies, and provide a conceptual basis for designing conformation-selective modulators in pain and inflammation.

## Introduction

Ligand-gated ion channels (LGICs) are multimeric membrane proteins that mediate rapid electrical and chemical signaling, underpinning synaptic transmission, muscle contraction, inflammatory processes, cardiovascular responses, and immune response[1–3, 10, 11]. ATP-activated P2X receptors, with their minimalist trimeric architecture and well-defined intersubunit ATP-binding sites, provide a tractable model for dissecting allosteric gating mechanisms[6–9].

Despite detailed structural snapshots of closed and open states, the field has long faced a disconnect between classical kinetic models—often invoking abstract intermediates such as the “flip state”[12, 13]—and the underlying molecular transitions that generate them. This “flip state” proposed to reconcile rapid ligand binding with slower pore opening, has remained phenomenological, lacking clear structural or energetic grounding. The challenge is particularly acute in homomeric channels like P2×2, where structural symmetry seems at odds with observed functional asymmetry and negative cooperativity[7].

A key unresolved question is how local ligand occupancy reshapes the energetic landscape to direct gating transitions—allowing identical subunits to follow ordered, asymmetric pathways. In principle, symmetry should enforce equivalent behavior, but because each ATP binding site bridges two neighboring subunits, partial occupancy and interfacial geometry can introduce local asymmetries.

Here, we bridge this conceptual gap by integrating legacy outside-out patch-clamp recordings [13] with a recursive Bayesian inference framework (MacroIR), enabling rigorous kinetic modeling from time-averaged macroscopic currents. By explicitly coupling ligand binding to independent subunit rotations, and validating these models with molecular dynamics simulations of receptors bound to a single ligand, we reveal that ATP lowers the rotational barrier for one subunit at the interface, while raising it for its neighbor. This stepwise, directional mechanism explains both negative cooperativity and the mechanistic basis of the classical flip state, reframing allosteric regulation as the dynamic modulation of transition pathways rather than simple equilibrium shifts.

Our findings reveal that functional asymmetry can emerge from the geometry of ligand-induced coupling in a structurally symmetric assembly, establishing a general principle for dynamic control in multimeric protein machines.

## Methods

### Overview

This study integrates electrophysiological recordings, molecular dynamics (MD) simulations, and Bayesian inference to investigate the gating mechanism of the P2×2 receptor. We developed a series of kinetic models, partially informed by MD simulations of the closed state bound to a single ATP molecule, and tested their performance against experimental data using Bayesian model comparison. A central component of our approach is the MacroIR algorithm, which extends our previous work [14] to enable likelihood estimation from time-averaged macroscopic currents. Detailed mathematical derivations, model specifications (in compliance with Bayesian Analysis Reporting Guidelines), and additional experimental and computational protocols are provided in the Supplementary Information (see SI S1 for full technical details).

### Electrophysiological Data and Preprocessing

We re-analyzed outside-out patch recordings from our previous study [13], obtained from HEK293 cells expressing rat P2×2 receptors. The dataset includes macroscopic responses to ATP pulses (0.1–10 mM, 0.2 or 10 ms duration), interleaved with 10 ms applications of 1 mM MgATP. Current traces were segmented into epochs for baseline, activation, and deactivation, with each segment resampled using exponentially increasing time intervals to efficiently capture both fast and slow dynamics. Baseline noise was quantified using intervals of 13.7 ms and 0.02 ms. **A full description of segmentation criteria, interval selection, and the preprocessing workflow is provided in Supplementary Information Section S1.1**.

### Kinetic Model Design and Bayesian Comparison

We evaluated nine kinetic schemes, including seven previously established models [13, 15] and two newly proposed conformational models inspired by recent structural studies of P2X receptors [8, 9]. The new schemes explicitly link ATP binding to partial subunit rotations and introduce either symmetric or asymmetric coupling between binding and gating transitions.

Models were grouped into three categories: state-based, allosteric, or conformational. A complete description of each scheme—including rate expressions, subunit interactions, and conductance assignments—is provided in Supplementary Information Section S1.2.

*Note: Two additional conformational models (previously referred to as Schemes VIII and XI) were excluded from the main analysis after preliminary assessment indicated they did not improve model fit or alter mechanistic conclusions. Full definitions and analysis of all eleven schemes are provided in the accompanying code and data repository*.

### Recursive Likelihood Estimation with MacroIR

All experimental measurements of ionic current, no matter how brief the integration time, are inescapably averages over finite intervals—there is no way to record a truly instantaneous current. As a result, time-averaging (whether due to low-pass filtering, hardware integration, or software binning) inevitably induces temporal correlations in the data, violating the assumptions of instantaneous observation and conditional independence that underlie traditional likelihood methods.

To address this, we developed the Macroscopic Interval Recursive (MacroIR) algorithm, a recursive Bayesian framework for inferring kinetic parameters from interval-averaged current data. MacroIR propagates the ensemble mean and covariance of channel state occupancies across arbitrarily long and non-uniform intervals, rigorously conditioning on the state at both the start and end of each interval. By leveraging these “boundary states,” MacroIR accurately captures the full statistics of each measurement window, without the need to enumerate all possible underlying state paths.

This boundary-conditioned approach contrasts with previous methods (e.g., Munch et al. [16]), which approximate the likelihood for each interval using only a single-point estimate (typically the mean current and variance at the interval midpoint or average). Such single-point approximations are practical for very short intervals but become unreliable when measurement intervals exceed the timescales of channel kinetics—as in exponentially scaled binning schemes—resulting in biased parameter estimates and underestimation of model evidence.

For comparison, we also implemented a non-recursive alternative, Macroscopic Interval Non-Recursive (MacroINR), which performs single-pass updates without propagating boundary state information. MacroIR thus overcomes the key limitations of prior approaches, enabling accurate and efficient Bayesian inference for kinetic models of arbitrary complexity and across a broad range of timescales.

All mathematical details, code (including validation scripts and usage examples), and a full implementation of MacroIR are provided in our public repository (https://github.com/lmoffatt/macrodrsubmission). Detailed derivations and algorithmic steps are presented in Supplementary Information Section S1.3.

### Molecular Dynamics Simulations

We investigated the structural basis of asymmetric gating in P2X receptors using molecular dynamics (MD) simulations. Models were constructed for the zebrafish P2X_4_ closed and open states (PDB: 3I5D and 4DW1), with added or modeled residues to ensure full-length chains and correct symmetry. An ATP molecule was docked into the A–B binding pocket of the closed state to create a singly liganded model. Simulations were run in explicit solvent using AMBER, followed by domain-wise analysis of conformational displacements projected onto closed → open transition vectors for key extracellular regions (head loop, LF, DF, LB). Details of system preparation, simulation protocols, and vector analysis are provided in Supplementary Information, Section S1.5.

## Data Availability

Processed experimental recordings and MCMC posterior samples are publicly available via Zenodo (https://doi.org/10.5281/zenodo.15085038) and GitHub (https://github.com/lmoffatt/macrodrsubmission/experiments). MD simulations inputs for AMBER and scripts for data analysis are also available in GitHub (https://gitlab.com/CLPF/p2x).

## Code Availability

Custom code for kinetic modeling, including all 9 schemes and 2 schemes not presented here, MacroIR/MacroINR likelihoods, and MCMC post-processing scripts, is available at https://github.com/lmoffatt/macro dr submission. The core engine is implemented in C++20; MCMC analysis and figure generation use RMarkdown.

### Bayesian Selection of Kinetic Schemes for P2X_2_ Activation

We used Bayesian model evidence to compare nine mechanistic schemes for ATP-induced activation of P2X_2_ receptors (Fig. 1), spanning classical state models, allosteric models, and subunit-resolved conformational mechanisms. Model evaluation was performed using outside-out patch-clamp recordings from rat P2X_2_ receptors stimulated with 0.2 ms ATP pulses (0.1–10 mM)[13]. Inference was carried out via affine-invariant parallel tempering MCMC within our recursive MacroIR framework, which rigorously accounts for temporal correlations in macroscopic current data. As a control, we also applied a non-recursive method (MacroINR) that is susceptible to temporal correlation errors (Supplementary Table S1).

The candidate schemes were grouped as follows:

- **State models:** Discrete closed, flipped, and open states.
- **Allosteric models:** Transition rate modulation without explicit structural mapping.
- **Conformational models:** ATP binding coupled to independent subunit rotations at the binding interface.

A conformational scheme with asymmetric subunit coupling (Scheme IX) was decisively favored, exceeding its symmetric counterpart (Scheme VIII) by a Bayes factor *>* 5,000 (Fig. 1d). The top-ranked non-conformational alternative (the subunit-specific allosteric Scheme VI) was disfavored by a factor of 6.4 (90% equal-tail interval: 5.5–6.7). Minimal state models and concerted allosteric schemes performed poorly (Extended Data Fig. 1).

These findings establish that:

1. Subunit rotation is the principal conformational step in activation;
2. ATP binding allosterically modulates rotational propensity at the interface;
3. This modulation is strongly asymmetric between subunits;
4. Macroscopic conductance scales with the number of rotated subunits.

Notably, the classical “flip state” is reinterpreted here as a probabilistic manifestation of underlying subunit transitions.

Model ranking was sensitive to likelihood approximation: the control method (MacroINR) systematically underestimated evidence for schemes with conformational intermediates, underscoring the importance of rigorous handling of temporal correlations (Figs. 1d–e).

#### Computational considerations

Bayesian model selection at this mechanistic depth is computationally intensive, with each scheme requiring 12–52 days to converge on a 16–32-core cluster. Explicitly convolving the patch-clamp amplifier’s low-pass filter kernel (10,kHz Bessel; 50,kHz digitization[17]) would increase computational demands more than tenfold, rendering such analyses impractical at present.

In summary, Bayesian evidence identifies a directional, asymmetrically coupled gating mechanism for P2X_2_, motivating further structural and functional investigation of this symmetry breaking.

### Posterior Analysis of Scheme IX

The posterior distributions of the parameters in Scheme IX (Fig. 2) fall into three distinct classes, reflecting how different aspects of the data constrain the model:

#### 1. Well-identified kinetic and noise parameters

Core parameters governing ligand binding (*b*_on_, *b*_off_), subunit rotation (*r*_on_, *r*_off_), channel conductance (), baseline current (γ), inactivation (*k*_inact_), channel number (*N*_ch_), and both white and pink noise (*ε* ^2^, *ν*^2^) all exhibit narrow, unimodal posterior distributions (posterior concentration factors (PCF) ranging from 16 to 68; Supplementary Table S4). This demonstrates that the 0.2 ms ATP-pulse data provide strong constraints on the fundamental kinetic rates and noise sources governing channel activation and deactivation.

#### 2. Moderately informed conductance scaling

Parameters describing how subunit rotation affects current—the closed-channel leakage ratio (*ρ*_leak_) and the rotation-conductance factor (*Rγ*)—show reproducible shifts from their priors (PCFs of 1.5 and 3, respectively), but retain appreciable uncertainty. The data are sensitive to the principal features of the current–rotation relationship, but do not fully resolve very low levels of resting (leak) current.

**Fig. 1.**
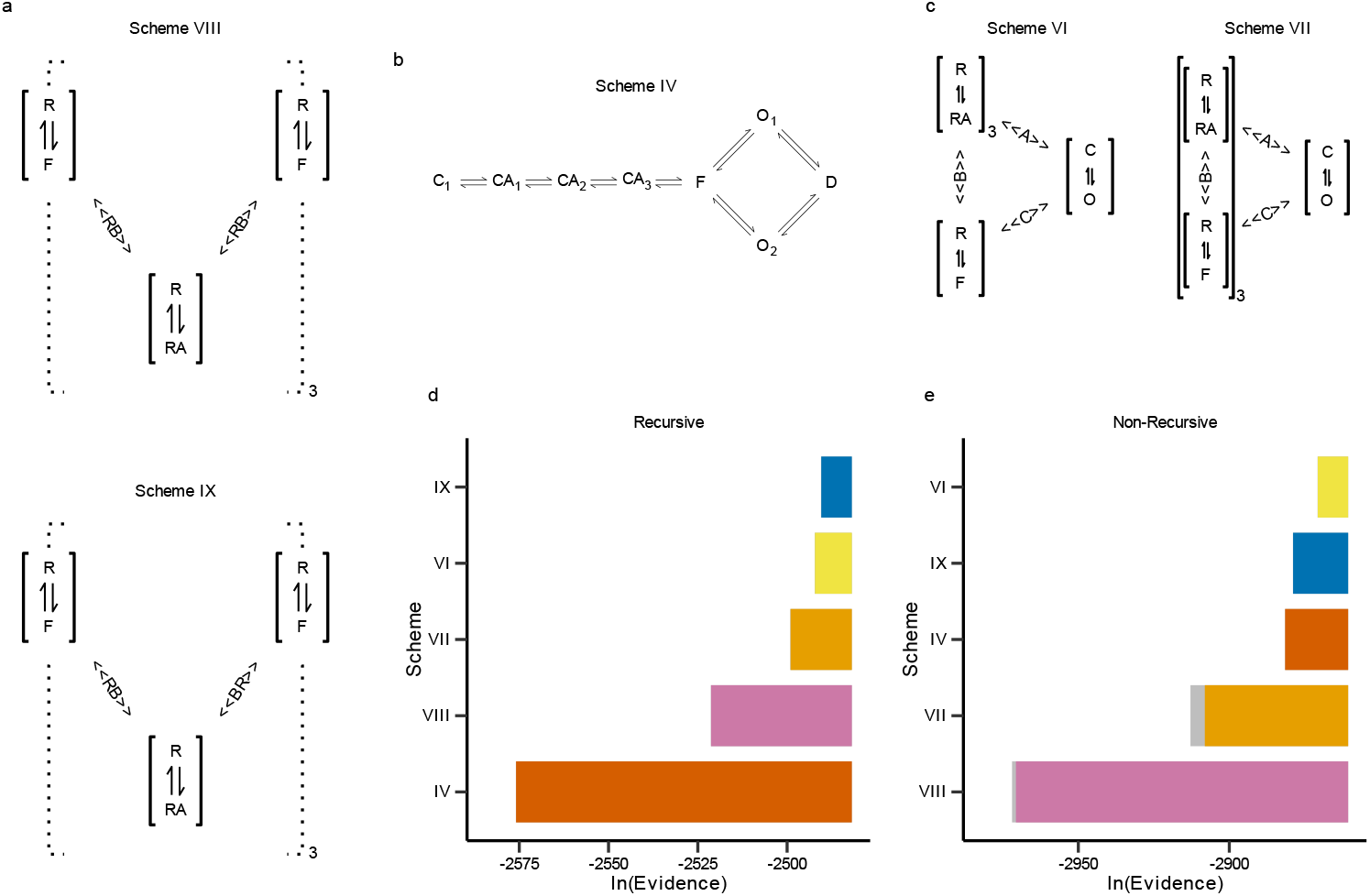
Bayesian evidence for conformational, allosteric, and state models. (**a**) Conformational models based on subunit rotation. R: resting; F: flipped; RA: agonist-bound; RB, BR: allosteric coupling via left or right subunits. (**b**) State model (Scheme IV) with a flip state branching into two open states. (**c**) Allosteric Schemes VI and VII, featuring concerted vs. subunit-specific flipping. (**d–e**) Bayesian evidence for each scheme computed with the recursive MacroIR algorithm (**d**) and the non-recursive MacroINR variant (**e**). Colors indicate model class; error bars denote standard errors of log_*e*_ (Evidence).

#### 3. Left-right ambiguity in binding-rotation coupling

Posterior distributions for the coupling constants that link ATP binding to subunit rotation (and vice versa) are distinctly bimodal (see Fig. 2 panels e, j, k). This bimodality arises because macroscopic currents alone cannot distinguish which subunit—at a given binding pocket—receives the stronger allosteric effect, so the MCMC sampler explores both mirror solutions. Although weakly informative priors can bias the sampler, the kinetic data preserve this ambiguity, reflecting the underlying symmetry of the homotrimer.

To resolve this degeneracy, we leveraged single-ATP molecular dynamics (MD) simulations (Section 2). These revealed that, upon ATP binding at an interface, the LF and Head domains of chain A and the DF domain of chain B move furthest toward the open conformation (Fig. 3, Table 1). Consistent with these structural displacements, we relabeled the posterior samples so that the stronger coupling (*RB*) is always assigned to chain A, and the weaker (*BR*) to chain B. This symmetry-breaking transformation collapses the bimodal posteriors into interpretable, unimodal distributions.

**Extended Data Fig. 1.**
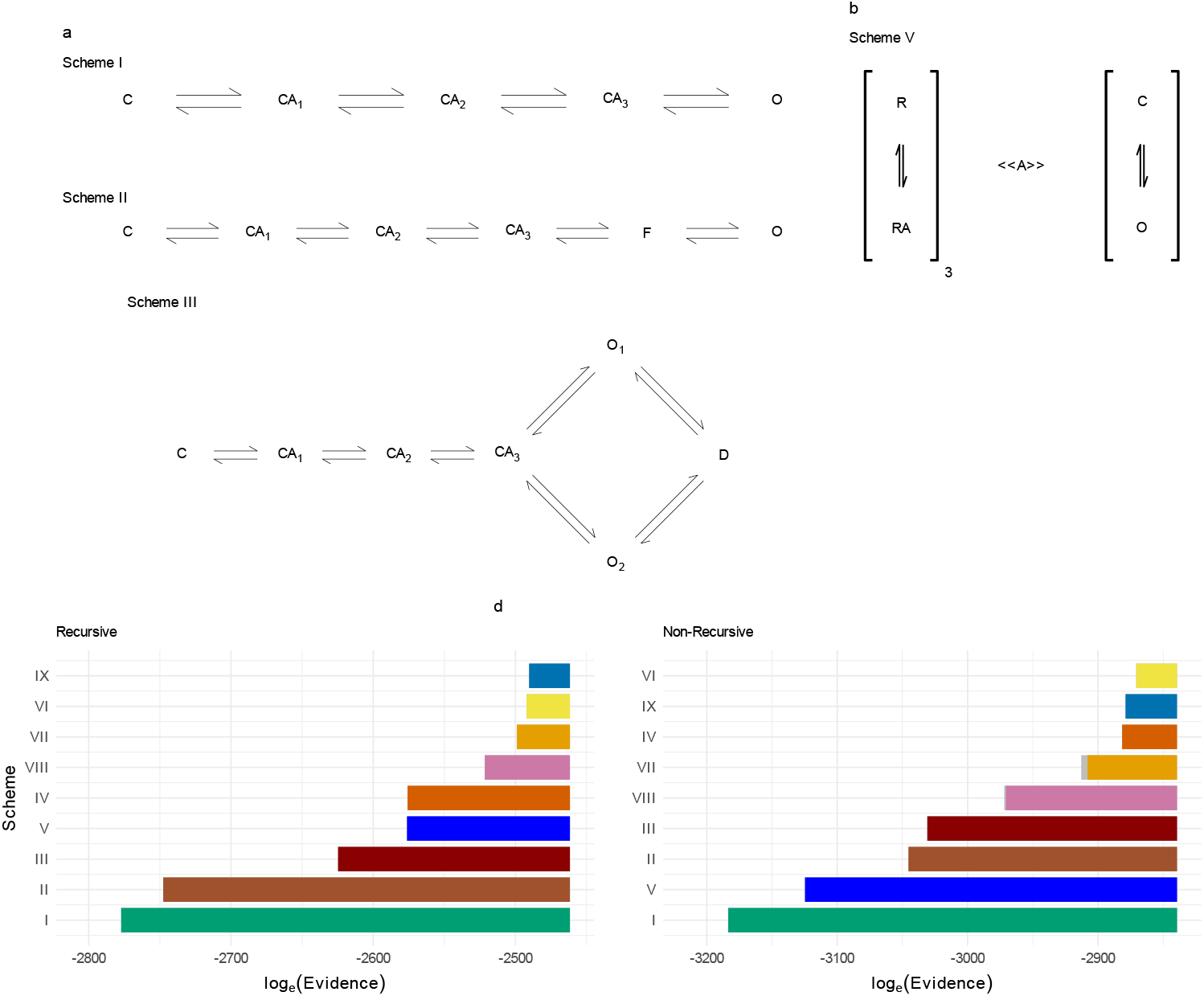
Expanded comparison of kinetic models. (**a**) Minimal state-based schemes (Schemes I–III). (**b**) Allosteric model Scheme V. (**c–d**) Bayesian evidence computed using MacroIR (**c**) and MacroINR (**d**). Colors and error bars as in Fig. 1.

Together, these three posterior patterns validate the kinetic model’s internal consistency, highlight which parameters are most strongly constrained by experiment, and set the stage for integrating kinetic and structural analyses to explain the functional basis of gating asymmetry.

### Single-bound Molecular Dynamics simulations clarify coupling asymmetry

To resolve the ambiguity in the binding–rotation coupling parameters identified by kinetic modeling, we analyzed Molecular Dynamics (MD) simulations of a closed-state P2X receptor with ATP bound at a single interface (chains A and B). For each structure in the ensemble, we quantified its Degree of Closed-to-Open Transition (DCOT) and its alignment with the opening direction (Degree of Closed-to-Open Direction, DCOD).

**Fig. 2.**
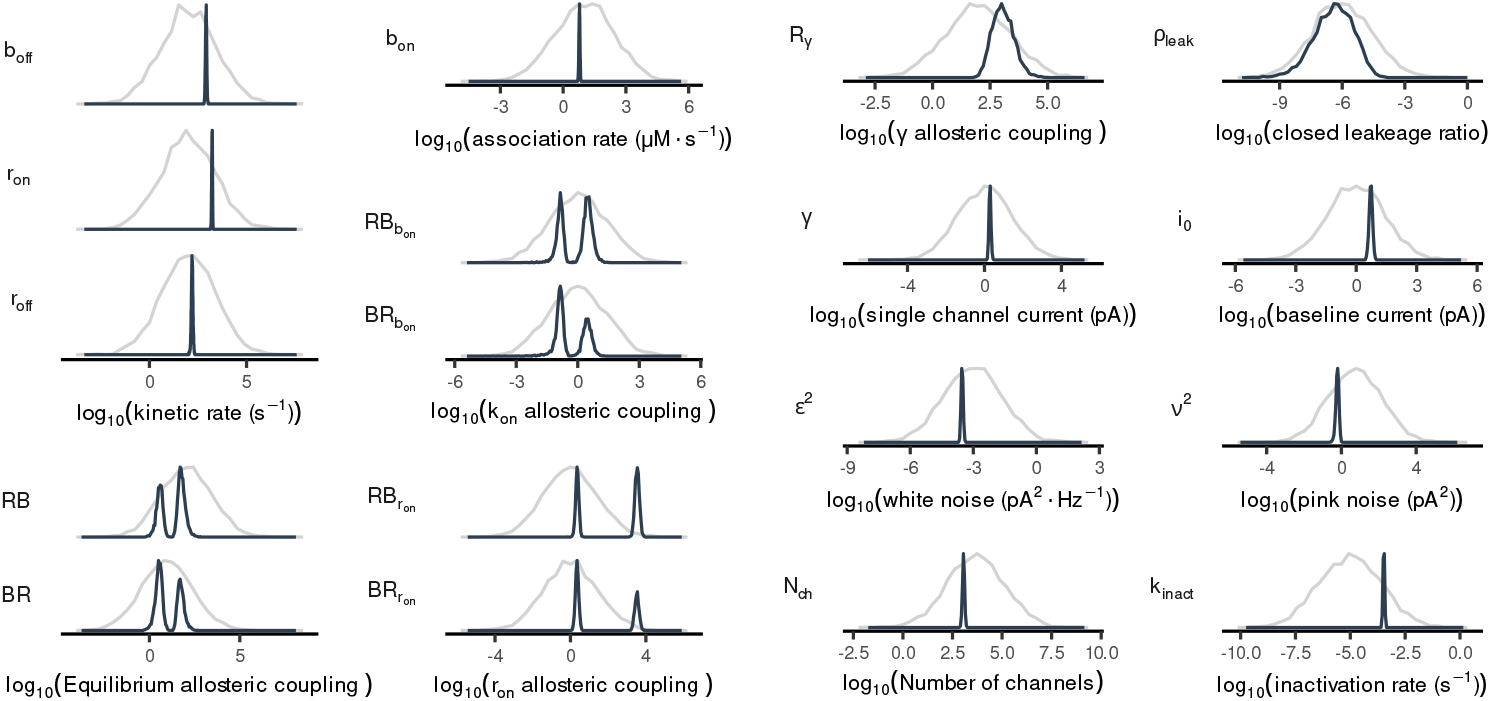
Posterior distributions of model parameters for Scheme IX. (**a**) Kinetic rate constants: *r*_on_ (rotation), *r*_off_ (return to resting state), and *b*_off_ (unbinding). (**b**) Association rate: *b*_on_ (binding). (**c**) Rotation-conductance coupling factor (*Rγ*). (**d**) Closed-channel leakage ratio (*ρ*_leak_). (**e**) Allosteric coupling factor affecting the binding rate 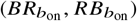. (**f**) Single-channel current of the fully rotated channel (*γ*). (**g**) Baseline current (*i*_0_). (**h**) White noise power (*ε* ^2^). (**i**) Pink noise power (*ν*^2^). (**j**) Equilibrium allosteric coupling factors (*BR, RB*). (**k**) Allosteric coupling factors affecting the rotation rate 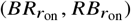. (**l**) Number of active channels (*N*_ch_). (**m**) Inactivation rate (*k*_inact_). The gray line indicates the prior; the colored line shows the posterior for each parameter. Probability densities are normalized to their maximum for comparability.

Figure 3 presents the DCOT analysis revealing the distinct behavioral patterns between the chains A and B. Among the four regions analyzed, the results for Dorsal Fin (DF) of chain B may initially seem counterintuitive, as the DCOT values shift opposite to the expected closed→open direction. This region forms part of the ATP-occupied binding cleft in 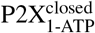, and its movement is strongly correlated with the Left Flipper (LF) of chain A [18, 19]. Notably, LF(A) exhibits a distribution markedly shifted toward the open conformation. This shift indicates a substantial alteration in its interaction pattern with DF(B), effectively triggering an initial “release” of DF(B) from the ATP-bound cleft and promoting movement in the direction opposite to the typical closed→open transition. In addition to the shift in LF distributions favoring chain A toward conformations closer to the open state, the Head-loop and Lower Body (LB) regions further emphasize this asymmetry, with chain A consistently adopting more open-like conformations. Particularly, the distinct displacements observed in the LB regions of both chains—which are structurally connected to the transmembrane domains[8]—suggest an initial asymmetrical channel activation mechanism triggered by the first ATP-binding event.

On the other hand, DCOD analysis (see Table 1) reveals that all examined-regions exhibit substantial movement in the closed-to-open direction, regardless of their interaction with ATP. The only exception is the Head-loop of chain B, which shows minimal movement along this pathway direction. This is in line with the fact that the Head domain plays a role in ATP recruitment but is not inherently coupled to the channel opening motion [20, 21]. However, when the Head-loop interacts with ATP (as in chain A), its movement shifts markedly toward the open conformation.

A similar analysis was conducted for chain C, with results available in Extended Data (Fig. 2, DCOT analysis) and in the Supplementary Information (Table **??**). Overall, the DCOT/DCOD results of chain C regions closely resemble those of the corresponding ones in chains A or B that are not interacting with ATP (Head-loop(C) aligns with Head-loop(B), and DF(C) with DF(A), etc).

This DCOT/DCOD analysis indicates a pronounced structural asymmetry in the receptor’s initial response to ATP binding, consistent with asymmetric coupling inferred from the kinetic analysis. By comparing MD and kinetic data, we assigned the stronger binding–rotation coupling (RB) to chain A and the weaker coupling (BR) to chain B. This structurally informed relabelling resolved the kinetic model’s symmetric degeneracy, collapsing bimodal parameter distributions into interpretable unimodal forms (Extended Data Fig. S2). In summary, MD simulations validate and clarify the asymmetric allosteric modulation indicated by Bayesian inference, directly linking structural rearrangements to the kinetic asymmetry observed in macroscopic currents. These findings set the stage for a detailed mechanistic interpretation of barrier modulation in the next section.

**Fig. 3.**
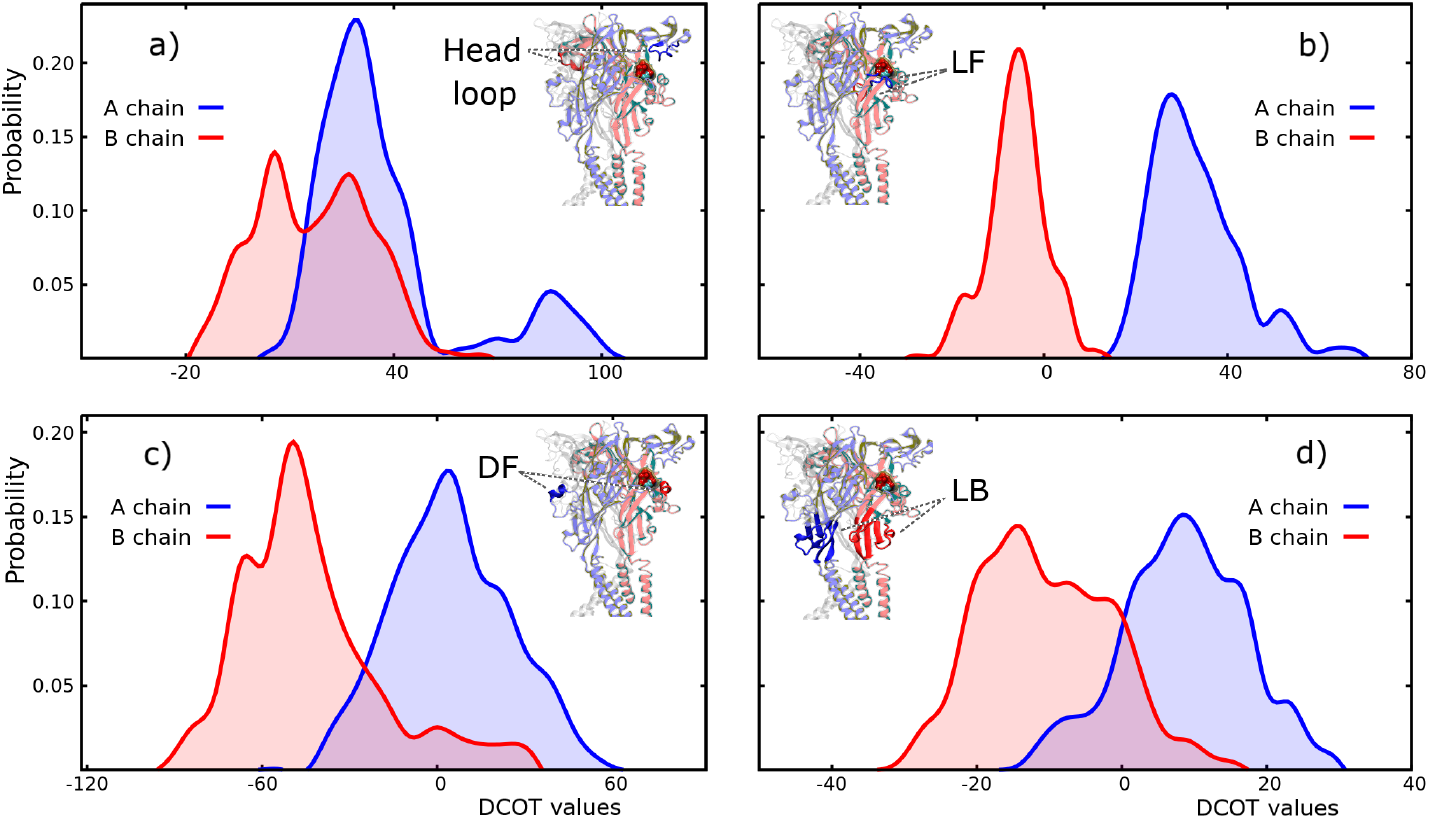
Normalized distribution of DCOT values (See Supplementary Information Eq. (S7)) of distinct regions of 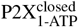. Chain A is shown in blue, and chain B is shown in red. Panel a) displays the results for the Head-Loop, b) for LF, c) for DF, and d) for LB. Each panel also includes a schematic illustration of the P2X receptor, with the specific region highlighted for clarity.

### Asymmetric Modulation of Transition Barriers by ATP Binding

To understand how ATP binding remodels the energetic landscape underlying P2X_2_ gating, we modeled three coupled transitions: ligand binding, rotation of the left subunit (subunit A at the A/B binding site), and rotation of the right subunit (subunit B at the same site) (Fig. 4a). Eachtransition was decomposed into *on* and *off* rate components, explicitly quantifying how ATP occupancy modulates both equilibrium and kinetic barriers. Conventional allosteric models encode coupling with a single equilibrium constant; here, we resolve modulation at the level of individual transition rates, allowing us to distinguish effects on both state stabilization and barrier height.

**Table 1.** DCOD averages (See Supplementary Information Eq. (S8)) of 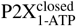 for LF, DF, LB and Head-Loop of chains A and B.

| Region | Chain | DCOD average | Standard deviation |
| --- | --- | --- | --- |
| Head-Loop | A | 46.04 | 22.12 |
| Head-Loop | B | 5.31 | 5.22 |
| LF | A | 85.83 | 16.16 |
| LF | B | 69.28 | 10.58 |
| DF | A | 72.19 | 19.45 |
| DF | B | 89.02 | 10.07 |
| LB | A | 76.07 | 10.01 |
| LB | B | 86.13 | 4.55 |

**Extended Data Fig. 2.**
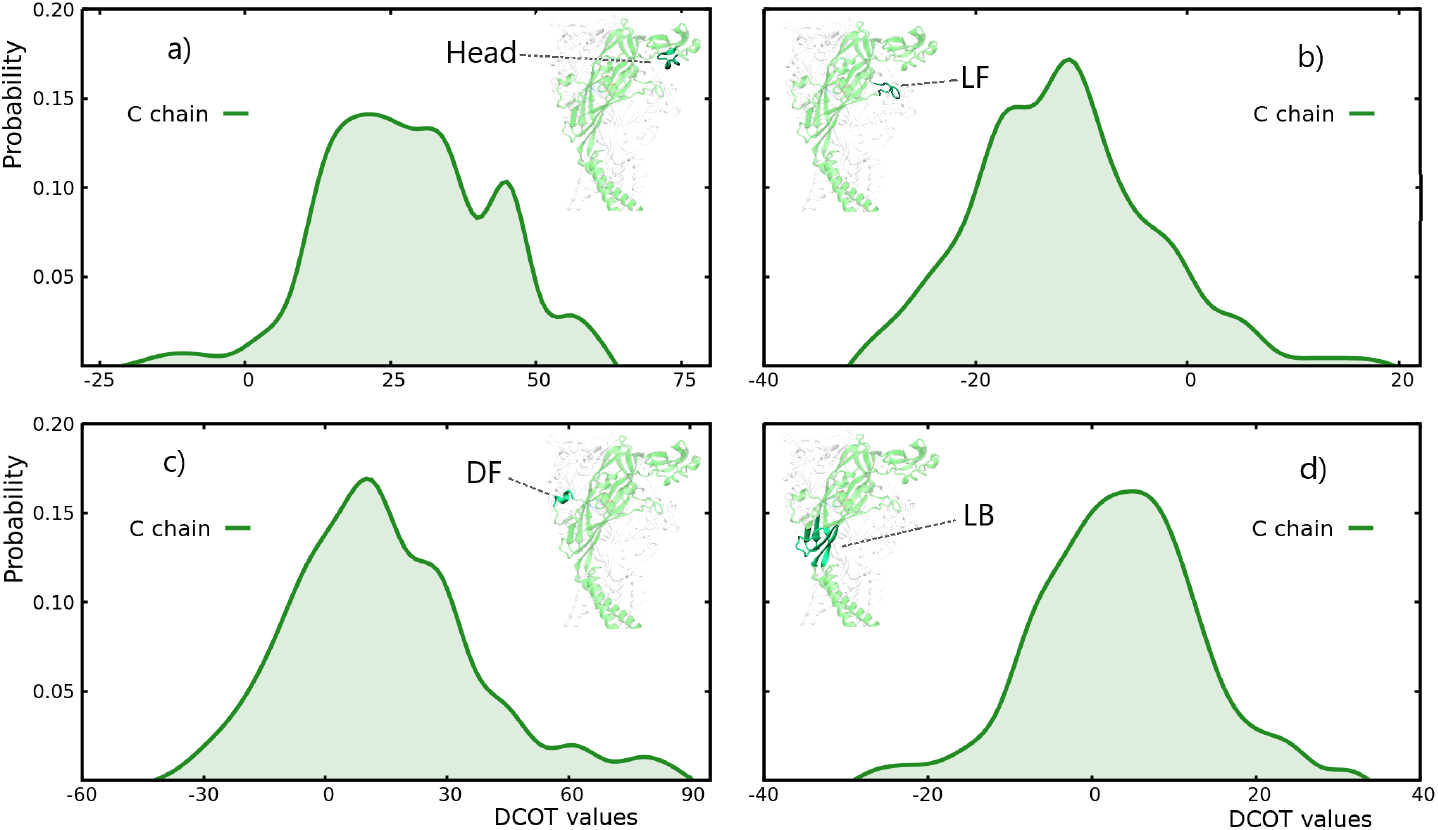
Normalized distribution of DCOT values (see Supplementary Information Eq. (S7)) for distinct regions of chain C in 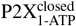. Panel a) shows the Head Loop, b) LF, c) DF, d) LB. Schematics depict the analyzed region on the P2X structure.

Posterior distributions (Fig. 4b,c) show that ATP binding strongly accelerates rotation of the left (A) subunit, but has minimal effect on the right (B) subunit. Similarly, rotation of subunit A substantially increases the rate of ATP binding 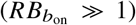, while rotation of subunit B has a weak or even inhibitory effect 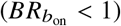. These patterns reveal a pronounced functional asymmetry in coupling, in agreement with both the kinetic and MD analyses.

A log–log analysis of coupling factors (Fig. 4d) further distinguishes between mechanistic regimes: (i) *regular allosterism* (stabilizing active states), (ii) *catalytic allosterism* (lowering activation barriers without a major equilibrium shift), and (iii) *inhibitory allosterism* (raising barriers to suppress transitions). ATP’s effect on subunit A exemplifies catalytic allosterism, enhancing both *on* and *off* rates, whereas BR coupling is weak, indicating minimal modulation of subunit B.

This asymmetric barrier tuning provides a mechanistic rationale for sequential, ordered subunit activation. By selectively lowering the barrier for rotation of subunit A, ATP binding directs the channel toward productive gating while limiting unproductive or energetically costly transitions. Thus, asymmetric allosteric modulation emerges as a general mechanism for controlling conformational flux and kinetic filtering in dynamic proteins.

In the following section, we quantitatively integrate these energetic insights into a full Markov model, connecting subunit-level transitions to macroscopic gating behavior.

### Integrating Subunit Conformational Dynamics into Channel Kinetics

To quantitatively link subunit-level conformational dynamics with macroscopic channel behavior, we constructed a Markov model explicitly representing all combinations of subunit binding and rotation states (Fig. 3). Each subunit undergoes ligand binding/unbinding and rotation/unrotation, with transition rates parameterized by the asymmetric allosteric couplings inferred previously (Fig. 4). Transition rates were scaled by the number of eligible subunits or sites, capturing both combinatorial multiplicity and asymmetric modulation.

This full kinetic framework enables calculation of state occupancies under both resting and stimulated conditions. Under ligand-free conditions (60 s post-ATP removal), the model predicts that stochastic fluctuations drive non-negligible occupancy of partially activated states: ∼ 15% of channels have one subunit rotated, and ∼ 1.1% have two (Fig. 5a). Thus, spontaneous baseline currents observed in patch-clamp experiments are quantitatively attributed to transitions into these partially active states, not to measurement noise. This prediction is confirmed by comparison to measured pre-ATP currents (see Fig. 6).

During ATP stimulation (10 mM pulses), state occupancies evolve sequentially: ligand binding precedes subunit rotation, yielding clear edge-to-center transition patterns (Fig. 5b). Deactivation, by contrast, proceeds via mixed ligand-bound and rotated states, highlighting distinct forward and reverse pathways.

Crucially, the model reveals that intermediate states are not merely transient: channels with two rotated subunits conduct 34 % of the maximal current (26–45 % equal-tailed interval; Fig. 5c) confirming that partial activation is physiologically significant—a result consistent with experimental findings that two occupied binding sites suffice for robust gating.

These results demonstrate that the observed spontaneous and partial activations are a direct consequence of the underlying Markovian kinetics. By tuning kinetic barriers, especially in the absence of ligand, the channel can minimize time spent in “risky” intermediate states, ensuring stability without sacrificing responsiveness. This provides a rigorous kinetic explanation for macroscopic currents observed in patch-clamp experiments and suggests that selective targeting of specific subunit transitions could provide new strategies for therapeutic modulation in P2X-related diseases.

In the following section, we directly validate the model’s predictive accuracy against experimental recordings.

**Extended Data Fig. 3.**
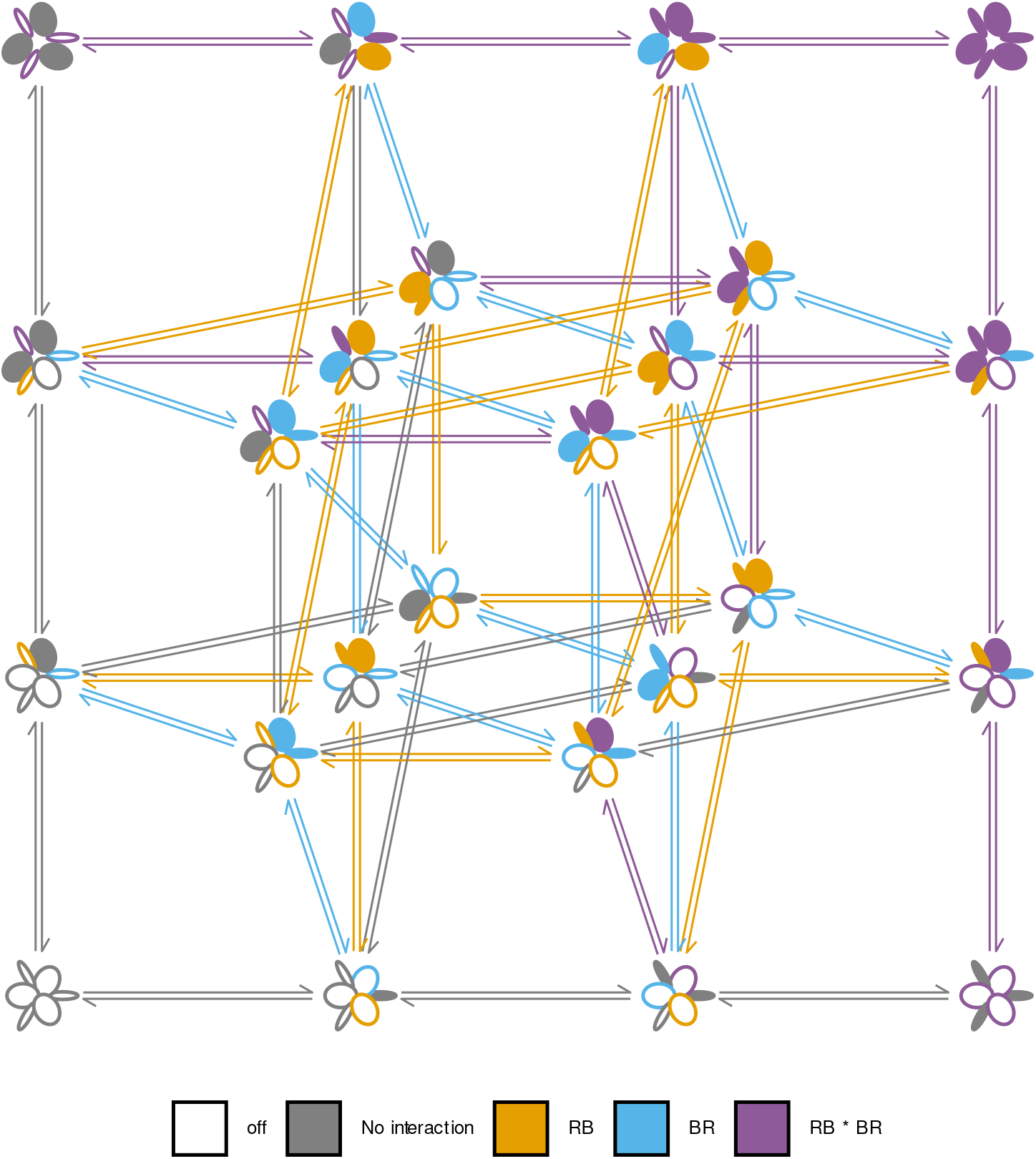
Kinetic Scheme IX: Comprehensive Representation of Subunit Interactions. Each subunit is depicted as an ellipse with nested binding sites; empty ellipses denote resting (unrotated/unbound) states, filled ellipses indicate activated (rotated/bound) states. Transition rates (vertical: rotation, horizontal: binding/unbinding) are color-coded: gray (baseline), orange (strong RB-type), light blue (moderate BR-type), purple (combined). The kinetic rate matrix is constructed from parameters in Fig. 4.

**Fig. 4.**
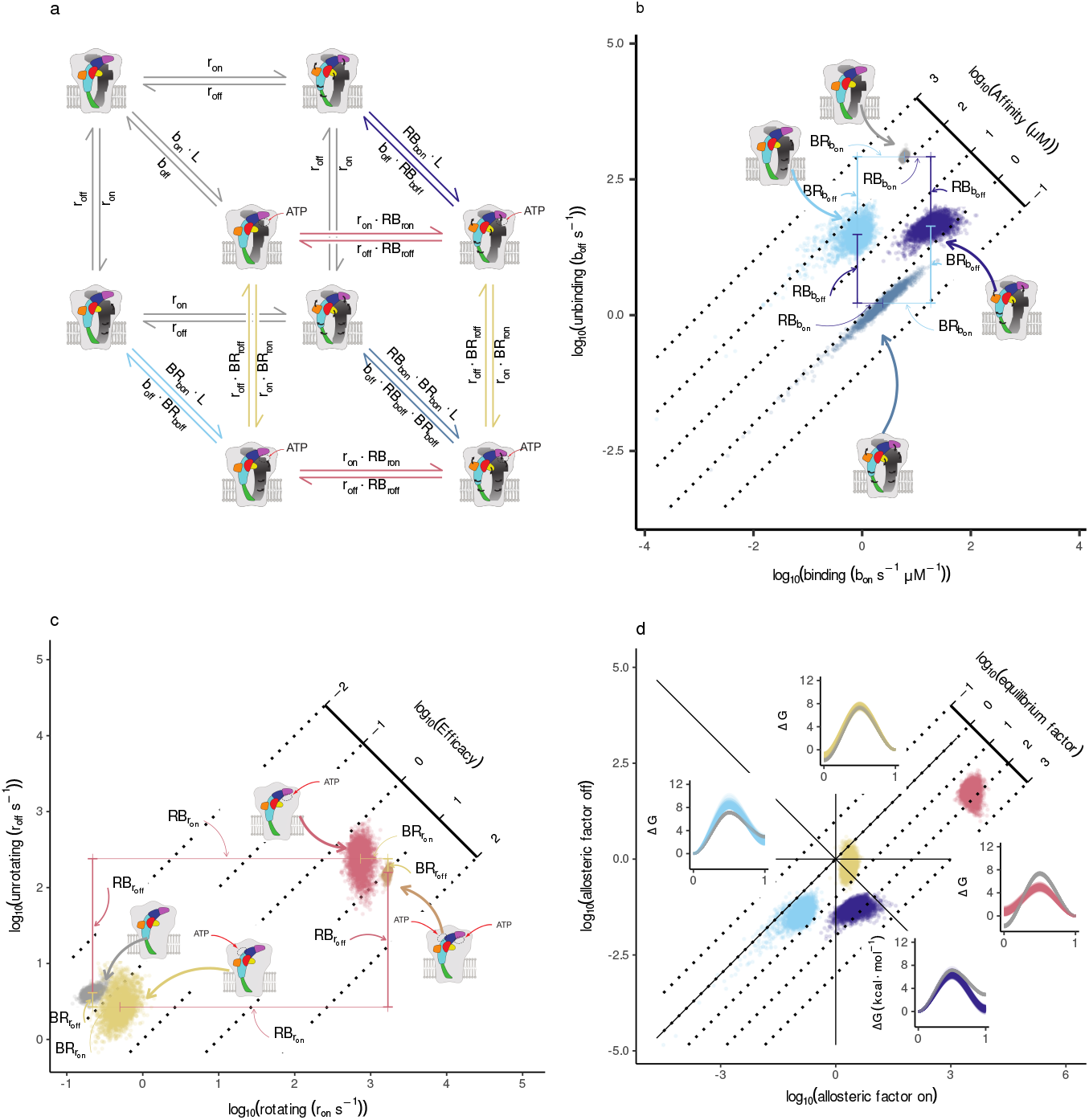
From Structure to Kinetic Barrier Modulation. (a) Cubic schematic illustrating three coupled transitions—agonist (ATP) binding, left subunit rotation, and right subunit rotation—where each vertex represents a distinct receptor state and each edge is labeled with its corresponding kinetic expression. Drawings adapted from [8] underscore the correspondence between model states and channel structures. Arrows indicate subunit rotations and ATP occupancy. (b, c) Posterior distributions for the rate constants of binding and rotation, segregating into four clusters that reflect the receptor’s structural states. (d) Log–log plot of the *on* versus *off* rate coupling factors, capturing how ATP binding or subunit rotations differentially accelerate or decelerate kinetic transitions. Insets illustrate the changes in energetic barriers, computed with a preexponential factor of 10^™6^ s.

### Validation of the Kinetic Model by Posterior Predictive Currents

To test the predictive capacity of our kinetic framework, we compared model-generated macroscopic currents with experimental recordings across a range of ATP concentrations. For each ATP pulse, MacroIR computes the expected current by integrating the state occupancies (Fig. 5a,b) with the unitary conductance of each state (Fig. 5c). Samples drawn from the posterior distribution closely reproduce both the amplitude and time course of the measured macroscopic response (Fig. 6a), validating the accuracy of the inferred channel kinetics.

**Fig. 5.**
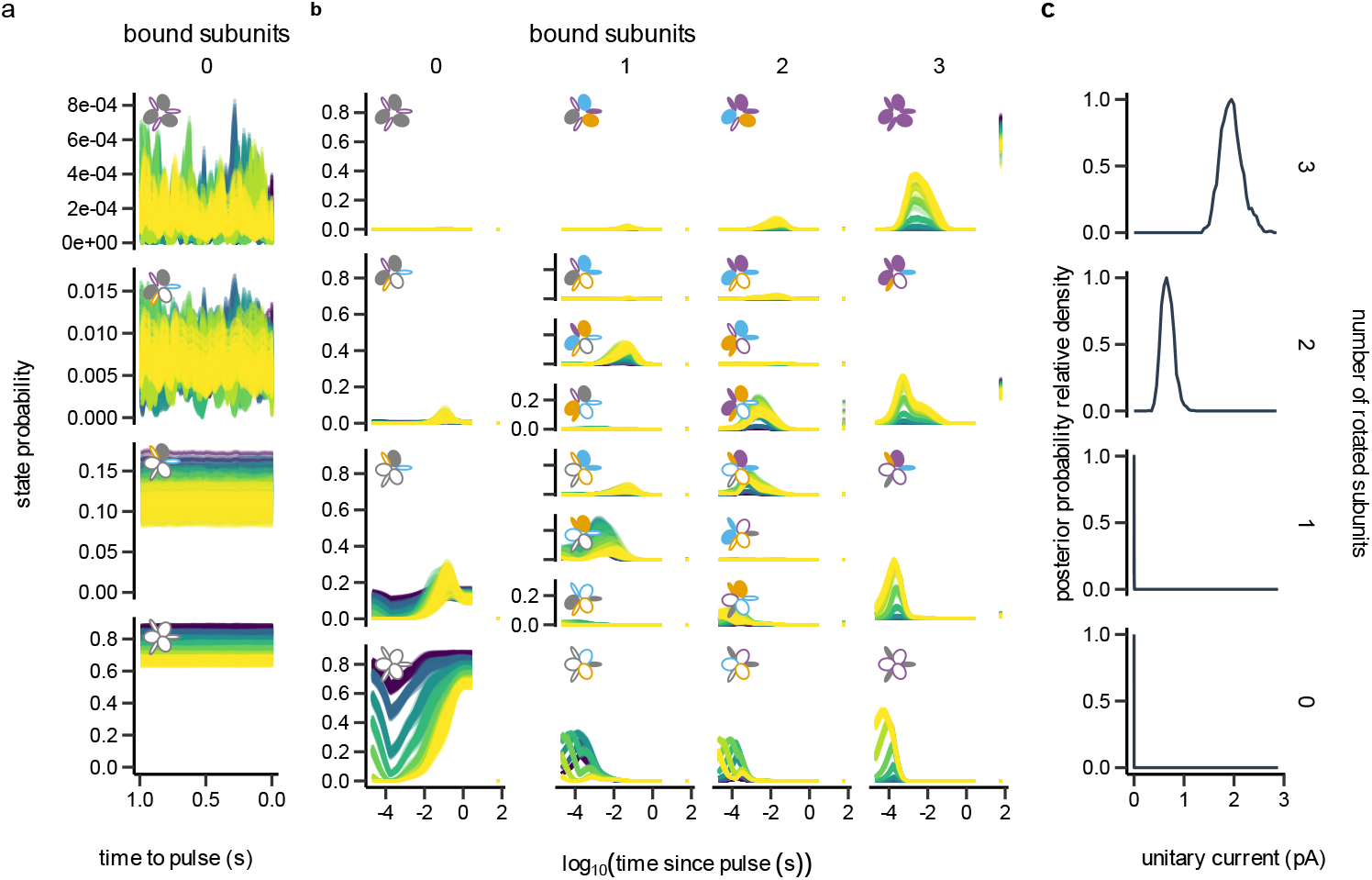
State Occupancies and Functional Currents Across Rotational States. (a) Equilibrium state occupancies 60 s after stimulus clearance, revealing significant pre-activation in the absence of ligand. (b) Temporal evolution of state probabilities during 10 mM ATP stimulation, showing sequential activation via ligand binding and subunit rotation. (c) Distribution of unitary currents across rotational states: channels with two rotated subunits conduct 34% (26–45 % equal-tailed interval) of maximal current, explaining baseline currents under ligand-free conditions.

Importantly, the model also quantitatively explains spontaneous current fluctuations observed prior to ATP application, attributing baseline activity to transitions into partially activated states rather than measurement noise alone.

MacroIR further partitions the total observed variance at each time point into three distinct components: (i) the variance arising from stochastic channel gating, modeled by the underlying Markov process; (ii) white noise, reflecting high-frequency, uncorrelated instrumental and thermal noise; and (iii) pink noise, representing slow, low-frequency baseline drift not captured by the channel model (Fig. 6b). During ATP pulses, the variance is dominated by the combination of modeled gating fluctuations and white noise, whereas pink noise becomes more prominent during the pre-pulse period.

Finally, direct comparison of the expected and observed likelihoods for each measurement (Fig. 6c) demonstrates excellent agreement between model and experiment, supporting both the validity of the probabilistic framework and the robustness of the parameter inference.

Collectively, these results show that MacroIR not only captures the dynamics and variability of ligand-gated channel activation and deactivation, but also provides a principled, quantitative explanation for spontaneous activity and macroscopic current fluctuations in the absence of ligand. This establishes MacroIR as a robust tool for mechanistic modeling and Bayesian inference in complex signaling systems.

**Fig. 6.**
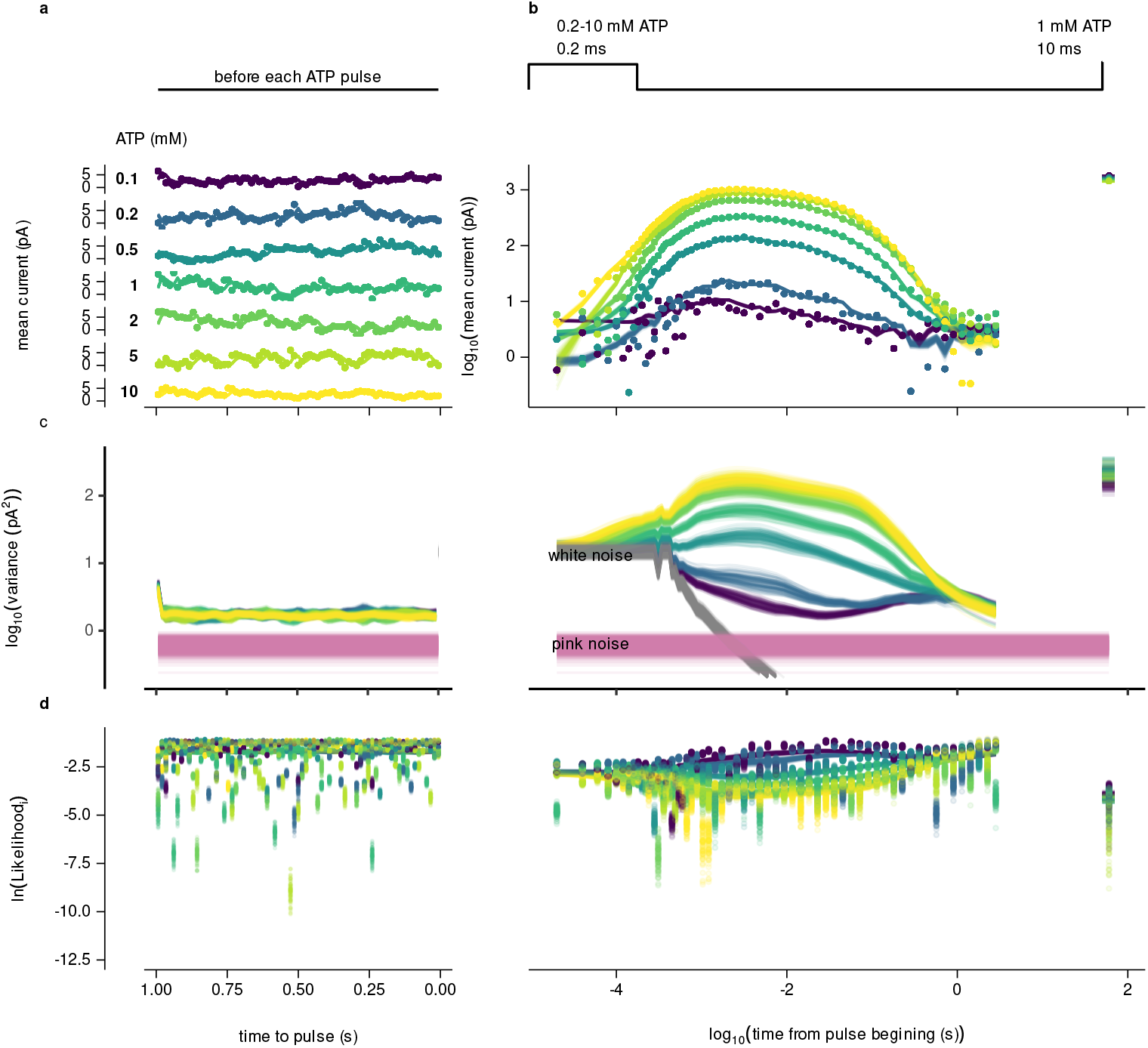
Predictive likelihood modeling from state dynamics: a posterior predictive check. (a) A sample from the posterior distribution of the predicted macroscopic current (blue) closely matches the measured current (black). (b) The total variance of the predicted current, with insets highlighting the contributions of white (red) and pink (green) noise. Across ATP concentrations (0.2 to 10 mM), white noise dominates during the ATP pulse, while pink noise is significant during the pre-pulse period. (c) Comparison of the expected (lines) and observed likelihoods, computed using a normal distribution for each measurement.

## Discussion

Despite its perfect threefold symmetry, the trimeric P2X receptor does not behave as a functionally equivalent trimer. We show that ATP binding at a single inter-subunit pocket selectively lowers the rotational barrier of one subunit while minimally affecting—and in fact slightly disfavoring—its neighbour. Once this first subunit rotates, the barrier for a second ATP to bind at the same interface rises, providing a direct mechanism for negative cooperativity and for the phenomenological “flip state” proposed in earlier kinetic schemes [7, 12, 22, 23]. Elevated barriers in the ligand-free receptor stabilise the closed conformation yet still permit rare excursions into partially rotated states, quantitatively accounting for spontaneous openings detected in outside-out patches.

Because ATP modulates *transition barriers* rather than simply shifting equilibria, substantial currents persist when only one or two sites are occupied. Channels with two rotated subunits conduct roughly half the maximal current, and the same kinetic scheme explains baseline activity without ligand. Such barrier modulation offers a design principle for dynamic proteins: by raising barriers in the unliganded state, the system suppresses wasteful transitions without altering equilibrium distributions, striking a balance between stability and responsiveness in a rugged energy landscape [24, 25].

Our favoured Scheme IX, containing 24 states dictated by first-principles coupling of binding to independent subunit rotation, is irreducible: removing any transition compromises quantitative fit or violates structural constraints. Bayesian comparison (Bayes factor *>* 5 000 against its symmetric counterpart) decisively selects this asymmetric, sequential mechanism over classical equilibrium models. Posterior analysis further groups parameters into well-identified kinetic and noise terms, moderately informed conductance scalings, and a left–right coupling degeneracy resolved by single-ATP molecular-dynamics simulations that independently revealed larger closed→open displacements in specific domains of chain A.

These mechanistic insights were possible only because MacroIR propagates ensemble moments across arbitrary integration windows, delivering an exact interval-averaged likelihood and closed-form log-evidence. Earlier approaches ignored gating noise [26, 27], relied on diagonal-covariance approximations [28], or idealised each sample as instantaneous [14, 29]. By completing this methodological arc, MacroIR allowed discrimination among subtle alternatives that lower-resolution analyses could not resolve, even though each model required 12–52 days on a 16–32-core cluster.

Ordered, asymmetric activation narrows the entropic search space, prevents simultaneous risky transitions, and yields graded conductance when not all binding sites are occupied—features likely advantageous for rapid synaptic signalling. Barrier-tuning may therefore govern other multimeric channels and allosteric machines in which partial agonism and negative cooperativity are common [30–33]. Mapping subunit-specific barriers thus opens a path to rational, subunit-selective modulators that tune gating dynamics rather than targeting static conformations [34, 35].

Scheme VI fitted almost as well as Scheme IX, and alternative topologies with additional microstates could plausibly capture nuances we did not test. All structural assignments derive from homology models and sub-microsecond simulations of the closed receptor; longer simulations and high-resolution structures of partially liganded states will be needed to confirm the inferred pathway. Although MacroIR rigorously explains spontaneous openings, we cannot exclude residual contributions from other channel populations or experimental artefacts. Single-channel recordings, targeted mutagenesis, and time-resolved cryo-EM will be essential to refine and generalise this kinetic-conformational framework.

By repositioning the flip state as an obligatory consequence of asymmetric barrier modulation, our work reframes allostery in dynamic proteins as control of transition *pathways* rather than static state populations. The integrative strategy presented here—combining high-precision electrophysiology, interval-exact Bayesian inference, and structurally informed simulations—provides a blueprint for uncovering latent mechanistic order in complex protein assemblies and for guiding mechanism-based therapeutics.

## Authors’ contributions

L.M. and G.P-S. conceived the study. L.M. performed the kinetic modeling and Bayesian inference. G.P-S. conducted the molecular dynamics simulations and structural analyses. Both authors discussed the results and contributed to the writing of the manuscript.

## Acknowledgments

We gratefully acknowledge the computational resources provided by CSC (Buenos Aires), UNC Supercómputo CCAD (Córdoba) and CCAR (Rosario); all part of SNCAD, Argentina. We thank the administrators and technical staff of these facilities for their support. L.M. and G.P. are members of CIC-CONICET. This work was supported by Universidad Nacional de Quilmes (PP827-942/20), CONICET and ANPCyT (ID:2436). The authors also thank Juliana Palma for helpful discussions and feedback.

## References

[1] Changeux, J. P., Devillers-Thiery, A. & Chemouilli, P. Acetylcholine receptor: an allosteric protein. Science 225 4668, 1335–45 (1984).

[2] Unwin, N. Neurotransmitter action: Opening of ligand-gated ion channels. Cell 72, 31–41 (1993).

[3] Lemoine, D. et al. Ligand-gated ion channels: New insights into neurological disorders and ligand recognition. Chem. Rev. 112, 6285–6318 (2012). PMID: 22988962.

[4] Galligan, J. Ligand-gated ion channels in the enteric nervous system. Neurogastroenterology & Motility 14, 611–623 (2002).

[5] Feske, S., Skolnik, E. Y. & Prakriya, M. Ion channels and transporters in lymphocyte function and immunity. Nature 12, 532–547 (2012).

[6] Nicke, A. et al. P2×1 and p2×3 receptors form stable trimers: a novel structural motif of ligand-gated ion channels. EMBO J. 17 (1998).

[7] Sattler, C. et al. Unravelling the intricate cooperativity of subunit gating in p2×2 ion channels. Sci. Rep. 10 (2020).

[8] Hattori, M. & Gouaux, E. Molecular mechanism of atp binding and ion channel activation in p2x receptors. Nature 485, 207–212 (2012).

[9] Kawate, T., Michel, J. C., Birdsong, W. T. & Gouaux, E. Crystal structure of the atp-gated p2×4 ion channel in the closed state. Nature 460, 592–598 (2009).

[10] Chessell, I. P. et al. Disruption of the p2×7 purinoceptor gene abolishes chronic inflammatory and neuropathic pain. Pain 114, 386–396 (2005).

[11] Fabre, J. et al. Decreased platelet aggregation, increased bleeding time and resistance to thromboembolism in p2y1-deficient mice. Nature 5, 1199–2012 (1999).

[12] Burzomato, V., Beato, M., Groot-Kormelink, P. J., Colquhoun, D. & Sivilotti, L. G. Single-channel behavior of heteromeric α1β glycine receptors: an attempt to detect a conformational change before the channel opens. J. Neurosci. 24, 10924–10940 (2004).

[13] Moffatt, L. & Hume, R. I. Responses of rat p2×2 receptors to ultrashort pulses of atp provide insights into atp binding and channel gating. J. Gen. Physiol. 130, 183–201 (2007).

[14] Moffatt, L. Estimation of ion channel kinetics from fluctuations of macroscopic currents. Biophys. J. 93, 74–91 (2007).

[15] Ding, S. & Sachs, F. Single channel properties of p2×2 purinoceptors. J. Gen. Physiol. 113, 695–720 (1999).

[16] Münch, J. L., Paul, F., Schmauder, R. & Benndorf, K. Bayesian inference of kinetic schemes for ion channels by kalman filtering. eLife 11, e62714 (2022).

[17] Qin, F., Auerbach, A. & Sachs, F. Hidden markov modeling for single channel kinetics with filtering and correlated noise. Biophys. J. 79, 1928–1944 (2000).

[18] Zhao, W.-S. et al. Relative motions between left flipper and dorsal fin domains favour p2×4 receptor activation. Nature 5, 4189 (2014).

[19] Pierdominici-Sottile, G., Moffatt, L. & Palma, J. The dynamic behavior of the p2×4 ion channel in the closed conformation. Biophys. J. 111, 2642–2650 (2016).

[20] Huang, L.-D. et al. Inherent dynamics of head domain correlates with atp-recognition of p2×4 receptors: Insights gained from molecular simulations. PLoS One 9, e97528 (2014).

[21] Racigh, V., Ormazábal, A., Palma, J. & Pierdominici-Sottile, G. Positively charged residues in the head domain of p2×4 receptors assist the binding of atp. J. Chem. Inf. Model. 60, 923–932 (2019).

[22] Jiang, R. et al. Intermediate closed channel state (s) precede (s) activation in the atp-gated p2×2 receptor. Channels 6, 398–402 (2012).

[23] Browne, L. E. & North, R. A. P2x receptor intermediate activation states have altered nucleotide selectivity. J. Neurosci. 33, 14801–14808 (2013).

[24] Frauenfelder, H., Sligar, S. G. & Wolynes, P. G. The energy landscapes and motions of proteins. Science 254, 1598–1603 (1991).

[25] Hilser, V. J. & Thompson, E. B. Intrinsic disorder as a mechanism to optimize allosteric coupling in proteins. Proc Natl Acad Sci USA 104, 8311–8315 (2007).

[26] Colquhoun, A. & Hawkes, A. G. On the stochastic properties of single ion channels. Philosophical Transactions of the Royal Society of London. B, Biological Sciences 300, 1–59 (1982).

[27] Sigworth, F. J. The variance of sodium current fluctuations at the node of ranvier. The Journal of Physiology 307, 97–129 (1980).

[28] Milescu, L. S., Akk, G. & Sachs, F. Maximum likelihood estimation of ion channel kinetics from macroscopic currents. Biophysical Journal 88, 2494–2515 (2005).

[29] Stepanyuk, A. R., Borisyuk, A. L. & Belan, P. V. Efficient maximum likelihood estimation of kinetic rate constants from macroscopic currents. Plos one 6, e29731 (2011).

[30] Changeux, J.-P. & Edelstein, S. J. Allosteric mechanisms in normal and pathological nicotinic acetylcholine receptors. Curr Opin Neurobiol 15, 369–377 (2005).

[31] Plested, A. J. R. Structural mechanisms of activation and desensitization in neurotransmitter-gated ion channels. Nat Struct Mol Biol 23, 494–502 (2016).

[32] Lee, C. H. et al. Nmda receptor structures reveal subunit arrangement and pore architecture. Nature 511, 191–197 (2014).

[33] Unwin, N. Refined structure of the nicotinic acetylcholine receptor at 4Å resolution. J Mol Biol 346, 967–989 (2005).

[34] North, R. A. P2x receptors as drug targets. Mol Pharmacol 90, 184–189 (2016).

[35] Burnstock, G. Purinergic signalling: pathophysiology and therapeutic potential. Pharmacological Reviews 63, 614–703 (2011).

