## Supplementary Information for "Bayesian inference of functional asymmetry in a ligand-gated ion channel"

### S1 Supplementary Methods

#### S1.1 Overview

This document provides a comprehensive description of all methodologies employed in this study, in full compliance with the Bayesian Analysis Reporting Guidelines (BARG). We detail:

- The experimental data and preprocessing used for kinetic modeling,
- The construction and conceptual basis of alternative kinetic schemes,
- The mathematical derivation and implementation of the likelihood function for time-averaged macroscopic currents,
- The MacroIR algorithm for recursive likelihood estimation,
- The computation of Bayesian evidence and procedures for posterior parameter inference and model comparison,
- And the structural characterization of the conformational ensemble of the closed P2X receptor—docked with an ATP molecule—sampled via molecular dynamics (MD) simulations.

Each subsequent section provides exhaustive technical details—including mathematical derivations, algorithmic implementation, and simulation protocols—to enable full reproducibility.

#### S1.2 Source and Preparation of P2X2 Macro-patch Data

**Data source and experimental context.** Kinetic models in this study were constrained using previously published macroscopic current recordings from outside-out patches expressing rat P2X2 receptors [1]. These high signal-to-noise datasets are ideally suited for kinetic modeling. The stimulation protocol comprised:

- 0.2 ms ATP pulses at varying concentrations (0.1, 0.2, 0.5, 1, 2, 10 mM) delivered every 2 minutes,
- Interleaved 10 ms ATP pulses at 1 mM to monitor response stability and estimate steady-state current,
- Precise timing between agonist delivery and current measurement, enabling accurate resolution of rapid activation events.

Comprehensive experimental details—including internal and external solutions, temperature, voltage protocol, and equipment—are provided in the original publication [1].

**Segmentation strategy for kinetic modeling.** To extract kinetic information across multiple timescales, raw current traces were segmented into defined intervals:

- *Pre-pulse baseline (1 s):* Divided into 72 intervals of 13.7 ms and 10 intervals of 0.02 ms to estimate baseline and pink noise.
- *Activation phase:* Raw (unaveraged) points were retained during each ATP pulse and for the 0.24 ms immediately preceding onset to resolve fast dynamics.
- *Deactivation phase:* Post-pulse decay was downsampled using exponentially increasing intervals, yielding approximately 10 data points per time decade.
- *Steady-state currents (10 ms pulses):* A single value was calculated by averaging the final half of each 10 ms pulse response, as a proxy for equilibrium current.

This multi-resolution preprocessing yielded the final dataset vectors used in the MacroIR-based Bayesian analysis.

#### S1.3 Kinetic Models Evaluated via Bayesian Comparison

Nine distinct kinetic models were evaluated via Bayesian inference (see Section **??**) to identify the most plausible gating mechanisms for P2X2 activation.

##### S1.3.1 Model Classes and Conceptual Overview

The models fall into three main categories:

- **State Models (Schemes I–IV):** Conventional Markov chains without explicit coupling, modeling transitions among closed, open, and flipped states [1, 2].
- **Allosteric Models (Schemes V–VII):** Incorporate phenomenological allosteric effects (e.g., MWC-type coupling between ligand binding and conformational change).
- **Conformational Models (Schemes VIII–IX):** Introduced in this study, these explicitly model subunit rotations and structurally guided allosteric interactions based on high-resolution P2X4 structures [3, 4].

##### S1.3.2 Previously Established Models (I–VII)

**State Models:** Include linear and branched schemes with closed, flipped, and open states. Scheme IV adds a bifurcated flip-to-open path to capture both fast and slow components of deactivation.

**Allosteric Models:** Schemes V–VII implement abstract coupling factors between binding, flipping, and opening transitions. Scheme VI assumes concerted flipping, whereas Scheme VII allows for subunit-specific flipping.

*Motivation for New Models:* While these models capture many features of the data, they lack structural specificity—particularly regarding the inter-subunit ATP-binding interface and the mechanical role of subunit rearrangements. These limitations motivated the development of mechanistically grounded conformational models.

##### S1.3.3 New Conformational Models (VIII–IX)

These two models replace the abstract flip state with ATP-induced subunit rotations. Binding sites reside at subunit interfaces, and coupling is defined between binding and adjacent subunit rotation (A = left, B = right). Coupling terms are:

- **RB:** Rotation(A)–Binding coupling.
- **BR:** Binding–Rotation(B) coupling.

The models differ in which interactions are enabled:

- **Scheme VIII:** RB = BR (symmetric coupling).
- **Scheme IX:** RB and BR distinct (asymmetric coupling).

##### S1.3.4 Parameterization of Conformational Models

**Conductance Submodel:** To avoid combinatorial explosion of open sub-states, conductance $i(n)$ was modeled as a saturating function of the number of rotated subunits $n$:

$$i(n) = i_{\text{max}} \cdot \frac{E_n}{E_n + 1}, \quad \text{with } E_n = E_0 \cdot F_g^n$$

where $E_0$ is basal efficacy and $F_g$ represents the per-rotation conductance gain.

**Kinetic Allosteric Coupling:** Coupling between transitions was implemented as multiplicative factors on the forward rate constants:

$$BR_{b_{\text{on}}} = \frac{b^*_{\text{on}}}{b_{\text{on}}}, \quad BR_{r_{\text{on}}} = \frac{r^*_{\text{on}}}{r_{\text{on}}}$$

where $b_{\text{on}}$ and $r_{\text{on}}$ are binding and rotation rates, respectively, and starred versions denote the rates when the allosteric partner is occupied.

#### S1.4 MacroIR: Recursive Bayesian Likelihood for Time-Averaged Currents

The MacroIR (Macroscopic Interval Recursive) algorithm is designed to compute the likelihood of time-averaged macroscopic current data for Bayesian inference on Markov models of ion channel kinetics. MacroIR propagates the mean and covariance of the state distribution over consecutive time intervals, allowing rigorous analysis of arbitrarily averaged or filtered current traces.

**Algorithm overview:** For a Markov model with $N$ states, transition rate matrix $\mathbf{Q}$, and state-wise current vector $\boldsymbol{\gamma}$, the probability of transitioning from state $i$ to $j$ in time $t$ is $P_{i\to j}(t) = [\exp(\mathbf{Q}t)]_{ij}$.

The expected current averaged over interval $t$ for paths starting in $i$ and ending in $j$ is:

$$\overline{\gamma}_{i\to j}(t) = \frac{1}{P_{i\to j}(t)} \sum_{k,n_1,n_2} V_{in_1} V^{-1}_{n_1 k} \gamma_k V_{kn_2} V^{-1}_{n_2 j} E_2(\lambda_{n_1} t, \lambda_{n_2} t) \tag{S1}$$

where $V$ is the eigenvector matrix of $\mathbf{Q}$, $\lambda_n$ are its eigenvalues, and $E_2(x,y) = (e^x - e^y)/(x - y)$ for $x \neq y$, $E_2(x,x) = e^x$.

The interval-averaged current mean and variance, as well as their propagation, are given by:

$$\overline{y}^{\text{pred}}_{0\to t} = N_{\text{ch}} \cdot (\boldsymbol{\mu}^{\text{prior}}_0 \cdot \overline{\boldsymbol{\gamma}}_0) \tag{S2}$$

$$\sigma^2_{\overline{y}^{\text{pred}}_{0\to t}} = \frac{\epsilon^2}{t} + \nu^2 + N_{\text{ch}}\left[\overline{\gamma}_0^T\left(\boldsymbol{\Sigma}_0^{\text{prior}} - \text{diag}(\boldsymbol{\mu}_0^{\text{prior}})\right)\overline{\gamma}_0 + \boldsymbol{\mu}_0^{\text{prior}}\cdot\mathbf{E}\right] \tag{S3}$$

where $\overline{\gamma}_0 = (E[\overline{\gamma}_{i\to\cdot}](t))_i$, and $\mathbf{E}_i = E[\overline{\gamma}^2_{i\to\cdot}](t)$.

The update and propagation steps for the mean and covariance of the state distribution are as follows:

$$\boldsymbol{\mu}_t^{\text{post}} = \boldsymbol{\mu}_0^{\text{prior}}\mathbf{P}(t) + \frac{\left(\overline{y}^{\text{obs}}_{0\to t} - \overline{y}^{\text{pred}}_{0\to t}\right)}{\sigma^2_{\overline{y}^{\text{pred}}_{0\to t}}}\boldsymbol{K}^{\mu}_{0\to t} \tag{S4}$$

$$\boldsymbol{\Sigma}_t^{\text{post}} = \boldsymbol{\Sigma}_t^{\text{prior}} - \frac{1}{\sigma^2_{\overline{y}^{\text{pred}}_{0\to t}}}(\boldsymbol{K}^{\mu}_{0\to t})^{\text{T}}\boldsymbol{K}^{\mu}_{0\to t} \tag{S5}$$

where $\mathbf{P}(t) = \exp(\mathbf{Q}t)$, and the Kalman gain-like term is:

$$\boldsymbol{K}^{\mu}_{0\to t} = \overline{\gamma}_0^T\left[\boldsymbol{\Sigma}_0^{\text{prior}} - \text{diag}\left(\boldsymbol{\mu}_0^{\text{prior}}\right)\right]\mathbf{P}(t) + \boldsymbol{\mu}_0^{\text{prior}}\cdot\left(\sum_j P_{i\to j}(t)\overline{\gamma}_{i\to j}(t)\right)_i\mathbf{P}(t) \tag{S6}$$

**Full implementation, validation notebooks, and usage examples are available at https://github.com/lmoffatt/macro_dr_submission. Additional mathematical details can be provided upon request.**

#### S1.5 Molecular Dynamics Simulations

##### S1.5.1 Model Construction

P2X receptors are homomeric or heteromeric assemblies of three monomers labeled $P2X_1$ through $P2X_7$. As of this writing, the Protein Data Bank includes experimental structures from various species for $P2X_1$ [5], $P2X_3$ [6–10], $P2X_4$ [3, 4, 11, 12], and $P2X_7$ [13–18]. Most available structures lack complete intracellular residues, which are essential for conducting stable MD simulations and preventing channel drift [19]. Furthermore, predictions with AlphaFold [20] for these regions typically have very low confidence.

Despite differences in permeation selectivity among P2X subtypes [21], the overall architecture—especially the extracellular region—is conserved. Each channel resembles a chalice, with each monomer having a dolphin-like shape and binding clefts located roughly 40 Å from the membrane surface. This conserved structure strongly suggests a unified closed→open pathway triggered by ATP binding. To date, $P2X_4$ and $P2X_7$ are the only subtypes with experimental structures available for both closed and open states [3, 4, 13, 15, 16].

To investigate ATP-induced perturbations, we generated three models of zebrafish $P2X_4$ (Uniprot F8W463): (1) a closed state, (2) an open state, and (3) a closed state bound to a single ATP molecule at the A-B interface ($P2X^{\text{closed}}_{\text{1-ATP}}$). Models (1) and (2) were based on PDB 3I5D (closed) and 4DW1 (open), respectively. Residue numbers were standardized by adding missing segments (Leu34, Asn35, Val360, Leu361) to chain-A, and residues 359-361 to chain-C, using chain-B as reference. In 4DW1, which contains only one monomer (Arg36-Ile359), missing residues were added and symmetry information used to reconstruct the trimer. Residues Val31, Gly32, and Thr33 were modeled with AlphaFold3 [20]. Thus, both $P2X^{closed}$ and $P2X^{open}$ span Val31–Leu361 in all three chains, though both lack the intracellular regions (Met1–Lys30, Thr362–Lys389).

For the single bound model, the ATP molecule was positioned at the A–B binding cleft of $P2X^{closed}$ by aligning the C$\alpha$ atoms of the open and closed conformations. Chain A contacts ATP via its head and Left Flipper (LF) domains; chain B via its Dorsal Fin (DF) domain. This construct is referred to as $P2X^{closed}_{1\text{-}ATP}$.

##### S1.5.2 Simulation Protocols

MD simulations were performed using the PMEMD module of AMBER24 [22]. Each model was solvated in an octahedral box of explicit TIP3P water molecules, with $Na^+$ and $Cl^-$ ions added for charge neutrality and 0.15 M ionic strength. Special care was taken to maintain internal water structure to preserve channel geometry during simulation.

Protein and water were described by the Amber19SB force field [23], while the POPC membrane (where relevant) used Lipid14 [24]. The simulation protocol was as follows:

1. Energy minimization (constant volume).
2. Gradual heating (NVT) from 0 K to 100 K over 500 ps.
3. Further heating (NPT) to 303 K. During both heating phases, $C_\alpha$ atoms were restrained with 1.5 kcal/mol Å$^2$.
4. Four consecutive equilibration runs of 10 ns each at 303 K, with harmonic restraints reduced stepwise: 0.5, 0.1, 0.05, 0.01 kcal/mol Å$^2$.
5. A final unrestrained equilibration of 200 ns.
6. Ten independent production runs of 200 ns each, initiated from the final equilibration snapshot with random Maxwellian velocities at 303 K. Coordinates were saved every 0.05 ns.

Electrostatic interactions were computed with the Particle Mesh Ewald method, applying a 10.0 Å cutoff radius. Thus, direct-space calculations were performed for $r < 10$ Å, while reciprocal-space calculations handled longer-range interactions [25, 26]. The SHAKE algorithm was applied to constrain hydrogen-involving bond lengths, enabling an integration time step of 2.0 fs. Stability of results was confirmed by comparing statistics from the first and second halves of each trajectory. **Note:** Complete simulation input files and data are available in the accompanying GitHub repository (https://gitlab.com/CLPF/p2x).

##### S1.5.3 Closed $\longrightarrow$ open transition vectors analysis

To analyze conformational changes, we defined vectors for key regions implicated in the P2X gating: LF (Lys285–Gly294), DF (Ser214–Cys220), LB (Asp59–Ser65, Asn195–Lys205, Arg261–Pro275, Phe327–Phe333), and the Head-Loop (Gly138–Thr149). Vectors describing the closed→open conformational change were defined for these regions. These vectors will be named as "transition vectors" and will be labeled as $LF^{trans}_{C->O}$, $DF^{trans}_{C->O}$, $LB^{trans}_{C->O}$, and Head-Loop$^{trans}_{C->O}$. To construct them, the open and closed structures were aligned, and for each region and monomer (A, B, C), the vector from closed to open ($V^{trans}_{C\to O}$) was computed by subtracting closed from open C$\alpha$ positions.

For each structure in the conformational ensemble of $P2X^{closed}_{1\text{-}ATP}$, we also computed vectors that describe the motion of each domain relative to the $P2X^{closed}$ conformation. These vectors will be refereed as "displacement vectors" and were built by subtracting the C$\alpha$ coordinates of

$P2X^{closed}$ from each MD snapshot (after alignment). Two quantitative metrics were computed for each region and conformation:

- **Degree of Closed-to-Open Transition (DCOT):**

$$\mathrm{DCOT} = 100 \times \frac{V_{C\to O}^{\text{trans}} \cdot V_i^{\text{disp}}}{|V_{C\to O}^{\text{trans}}|} \tag{S7}$$

  This measures the projection of the displacement onto the closed→open direction, normalized to the transition vector's length.

- **Degree of Closed-to-Open Direction (DCOD):**

$$\mathrm{DCOD} = 100 \times \frac{|V_{C\to O}^{\text{trans}} \cdot V_i^{\text{disp}}|^2}{|V_{C\to O}^{\text{disp}}|^2} \tag{S8}$$

  This quantifies the squared magnitude of the movement in the open direction, normalized to the displacement's length.

For both metrics, $V$ refers to any of the four studied-regions, and $i$ indexes all sampled conformations.

### References


[1] Moffatt, L. & Hume, R. I. Responses of rat p2x2 receptors to ultrashort pulses of atp provide insights into atp binding and channel gating. *J. Gen. Physiol.* **130**, 183–201 (2007).

[2] Ding, S. & Sachs, F. Single channel properties of p2x2 purinoceptors. *J. Gen. Physiol.* **113**, 695–720 (1999).

[3] Kawate, T., Michel, J. C., Birdsong, W. T. & Gouaux, E. Crystal structure of the atp-gated p2x4 ion channel in the closed state. *Nature* **460**, 592–598 (2009).

[4] Hattori, M. & Gouaux, E. Molecular mechanism of atp binding and ion channel activation in p2x receptors. *Nature* **485**, 207–212 (2012).

[5] Bennetts, F. M. *et al.* Structural insights into the human p2x1 receptor and ligand interactions. *Nature* **15**, 8418 (2024).

[6] Mansoor, S. E. *et al.* X-ray structures define human p2x3 receptor gating cycle and antagonist action. *Nature* **538(7623)**, 66–71 (2016).

[7] Wang, J. *et al.* Druggable negative allosteric site of p2x3 receptors. *Science* **115**, 4939–4944 (2018).

[8] Li, M. *et al.* Molecular mechanisms of human p2x3 receptor channel activation and modulation by divalent cation bound atp. *Elife* **8**, e47060 (2019).

[9] Thach, T. *et al.* Mechanistic insights into the selective targeting of p2x3 receptor by camlipixant antagonist. *Journal of Biological Chemistry* **301** (2025).

[10] Kim, G.-R. *et al.* Discovery of triazolopyrimidine derivatives as selective p2x3 receptor antagonists binding to an unprecedented allosteric site as evidenced by cryo-electron microscopy. *Journal of Medicinal Chemistry* **67**, 14443–14465 (2024).

[11] Shen, C. *et al.* Structural insights into the allosteric inhibition of p2x4 receptors. *Nature* **14**, 6437 (2023).

[12] Shi, H., Ditter, I. A., Oken, A. C. & Mansoor, S. E. Human p2x4 receptor gating is modulated by a stable cytoplasmic cap and a unique allosteric pocket. *Science* **11**, eadr3315 (2025).

[13] McCarthy, A. E., Yoshioka, C. & Mansoor, S. E. Full-length p2x7 structures reveal how palmitoylation prevents channel desensitization. *Cell* **179**, 659–670 (2019).

[14] Kasuya, G. *et al.* Structural insights into divalent cation modulations of atp-gated p2x receptor channels. *Cell Reports* **14**, 932 – 944 (2016).

[15] Karasawa, A. & Kawate, T. Structural basis for subtype-specific inhibition of the p2x7 receptor. *elife* **5**, e22153 (2016).

[16] Sheng, D. *et al.* Structural insights into the orthosteric inhibition of p2x receptors by non-atp analog antagonists. *Elife* **12**, RP92829 (2024).

[17] Oken, A. C. *et al.* High-affinity agonism at the p2x7 receptor is mediated by three residues outside the orthosteric pocket. *Nature* **15**, 6662 (2024).

[18] Oken, A. C. *et al.* P2x7 receptors exhibit at least three modes of allosteric antagonism. *Science* **10**, eado5084 (2024).

[19] Pierdominici-Sottile, G., Moffatt, L. & Palma, J. The dynamic behavior of the p2x4 ion channel in the closed conformation. *Biophys. J.* **111**, 2642–2650 (2016).

[20] Abramson, J. *et al.* Accurate structure prediction of biomolecular interactions with alphafold 3. *Nature* **630**, 493–500 (2024).

[21] Samways, D. S., Li, Z. & Egan, T. M. Principles and properties of ion flow in p2x receptors. *Frontiers in cellular neuroscience* **8** (2014).

[22] Case, D. *et al.* Amber 18. *University of California, San Francisco* (2018).

[23] Tian, C. *et al.* ff19sb: amino-acid-specific protein backbone parameters trained against quantum mechanics energy surfaces in solution. *Journal of chemical theory and computation* **16**, 528–552 (2019).

[24] Dickson, C. J. *et al.* Lipid14: The amber lipid force field. *Journal of Chemical Theory and Computation* **10**, 865–879 (2014).

[25] Darden, T., York, D. & Pedersen, L. Particle mesh ewald: An n.log(n) method for ewald sums in large systems. *The Journal of Chemical Physics* **98**, 10089–10092 (1993).

[26] Essmann, U. *et al.* A smooth particle mesh ewald method. *The Journal of Chemical Physics* **103**, 8577–8593 (1995).


### Supplementary Tables

| Al.[a] | Sch.[b] | Pr. [c] | T [d] | It. [e] | ESS [f] | ln(Ev) [g] | 90% CI [h] | $\hat{R}$ [i] | Converges[j]? |
|---|---|---|---|---|---|---|---|---|---|
| R | I | 16 O | 48 | 518 | 41,250 | −2,776.9 | (-2,777.0, -2,776.9) | 1 | ✓ |
| R | II | 16 O | 48 | 528 | 43,065 | −2,747.4 | (-2,747.4, -2,747.3) | 1.0001 | ✓ |
| R | III | 16 O | 47 | 482 | 34,778 | −2,624.6 | (-2,624.6, -2,624.5) | 1.0005 | ✓ |
| R | IV | 32 X | 13 | 773 | 28,964 | −2,575.7 | (-2,575.8, -2,575.7) | 1.0012 | ✓ |
| R | IV | 32 O | 45 | 399 | 10,118 | −2,575.7 | (-2,575.7, -2,575.6) | 1.0007 | ✓ |
| R | V | 16 O | 48 | 525 | 43,049 | −2,576.1 | (-2,576.1, -2,576.1) | 1 | ✓ |
| R | VI | 16 X | 47 | 655 | 10,219 | −2,492.1 | (-2,492.2, -2,492.1) | 1 | ✓ |
| R | VII | 16 X | 44 | 114 | 1,845 | −2,501.3 | (-2,501.4, -2,501.1) | 1.0112 | ✓ |
| R | VII | 32 E | 52 | 214 | 6,062 | −2,498.9 | (-2,499.0, -2,498.8) | 1.0062 | ✓ |
| R | VIII | 16 O | 44 | 184 | 7,901 | −2,516.9 | (-2,517.0, -2,516.9) | 1.0001 | ✓ |
| R | IX | 32 X | 12 | 260 | 15,988 | −2,521.3 | (-2,521.4, -2,521.3) | 1 | ✓ |
| R | IX | 16 O | 44 | 207 | 10,719 | −2,521.2 | (-2,521.2, -2,521.1) | 1 | ✓ |
| R | X | 32 X | 44 | 519 | 8,846 | −2,490.3 | (-2,490.4, -2,490.3) | 1.0005 | ✓ |
| R | XI | 32 X | 44 | 406 | 6,311 | −2,498.9 | (-2,498.9, -2,498.8) | 1.0002 | ✓ |
| NR | I | 16 O | 48 | 867 | 80,108 | −3,183.4 | (-3,183.4, -3,183.3) | 1 | ✓ |
| NR | II | 16 O | 48 | 778 | 66,764 | −3,045.1 | (-3,045.1, -3,045.1) | 1 | ✓ |
| NR | III | 16 O | 48 | 728 | 56,459 | −3,030.7 | (-3,030.8, -3,030.7) | 1.0028 | ✓ |
| NR | IV | 32 O | 45 | 519 | 35,859 | −2,881.3 | (-2,881.3, -2,881.2) | 1.0001 | ✓ |
| NR | V | 16 O | 48 | 734 | 58,425 | −3,124.5 | (-3,124.6, -3,124.5) | 1 | ✓ |
| NR | VI | 16 X | 47 | 735 | 11,848 | −2,870.6 | (-2,870.6, -2,870.5) | 1.0004 | ✓ |
| NR | VII | 16 X | 45 | 123 | 3 | −3,008.9 | (-3,014.0, -3,003.7) | 1.2094 | ✗ |
| NR | VII | 32 E | 48 | 231 | 4 | −2,910.4 | (-2,914.3, -2,906.5) | 1.1635 | ✣ |
| NR | VIII | 16 O | 44 | 259 | 3,582 | −2,956.4 | (-2,956.5, -2,956.3) | 1 | ✓ |
| NR | IX | 16 O | 45 | 376 | 52 | −2,971.1 | (-2,972.2, -2,970.1) | 1.0205 | ✓ |
| NR | X | 16 X | 48 | 466 | 24,935 | −2,878.7 | (-2,878.7, -2,878.6) | 1 | ✓ |
| NR | XI | 16 X | 47 | 441 | 7,123 | −2,893.1 | (-2,893.2, -2,893.0) | 1.0042 | ✓ |

**Table S1**: Convergence analysis of the Evidence Evaluation for different kinetic schemes and likelihood approximation algorithms. [a] Algorithm used (R: recursive; NR: non-recursive approximation). [b] Scheme identifier (I to XI). [c] Processor configuration (number of CPUs and model, e.g. O=AMD Opteron 6276, X=Xeon E5-2670, E=AMD EPYC 7302P). [d] Time duration of the Monte Carlo Markov Chain simulations in days. [e] Number of iterations (in thousands). [f] Effective sample size of the chain. [g] Natural logarithm of the Bayesian Evidence. [h] Equal Tail 90% credible interval for ln(Evidence). [i] Potential scale reduction factor ($\hat{R}$) used to assess convergence. [j] Convergence indicator (green check: converged; orange/red: close to convergence or non-convergence).

| Parameter[a] | Full Name[b] | Units[c] | DM[c] | Median[f] | gsd [g] | $90\% CI$[h] | $\hat{R}$ [i] | ESS [j] | $PCF_{sd}$ [k] | $PCF_{CI}$[l] |
|---|---|---|---|---|---|---|---|---|---|---|
| $b_{on}$ | Binding on | $\mu M^{-1}\, s^{-1}$ | | 6.02 | 1.06 | (5.45, 6.63) | 1.0002 | 5,200 | 54.7 | 54.4 |
| $b_{off}$ | Binding off | $s^{-1}$ | | 826 | 1.08 | (727, 935) | 0.9998 | 4,760 | 42.6 | 43.2 |
| $r_{on}$ | Rotation on | $s^{-1}$ | | 1,680 | 1.05 | (1,540, 1,810) | 1.0002 | 5,160 | 65.9 | 68 |
| $r_{off}$ | Rotation off | $s^{-1}$ | | 157 | 1.11 | (132, 185) | 1 | 4,930 | 30.9 | 31.1 |
| $RB$ | RB Equilibrium Coupling | | | 38.8 | 4.42 | (2.21, 125) | 1.0006 | 5,320 | 2.17 | 2.61 |
| $BR$ | BR Equilibrium Coupling | | | 5.11 | 4.41 | (1.66, 94.7) | 1.0003 | 5,360 | 2.19 | 2.63 |
| $RB_{r_{on}}$ | RB Rotation Coupling | | | 2,580 | 37.1 | (1.81, 4,990) | 1.0013 | 4,980 | 0.91 | 1.36 |
| $BR_{r_{on}}$ | BR Rotation Coupling | | | 2.76 | 36.9 | (1.7, 4,540) | 1.0012 | 5,030 | 0.92 | 1.39 |
| $RB_{b_{on}}$ | RB Binding Coupling | | | 1.92 | 5.35 | (0.08, 6.56) | 1.0002 | 5,040 | 1.95 | 2.42 |
| $BR_{b_{on}}$ | BR Binding Coupling | | | 0.19 | 5.42 | (0.06, 5.27) | 1.0006 | 5,390 | 1.93 | 2.39 |
| $RB$ | RB Equilibrium Coupling | | X | 56.5 | 1.61 | (30.6, 144) | 0.9999 | 5,470 | 5.88 | 5.87 |
| $BR$ | BR Equilibrium Coupling | | X | 3.49 | 1.57 | (1.49, 6.49) | 1 | 5,320 | 6.13 | 6.21 |
| $RB_{r_{on}}$ | RB Rotation Coupling | | X | 3,400 | 1.31 | (2,190, 5,300) | 1 | 5,110 | 12.2 | 12.1 |
| $BR_{r_{on}}$ | BR Rotation Coupling | | X | 2.26 | 1.22 | (1.61, 3.15) | 1.0005 | 4,870 | 16.5 | 16.4 |
| $RB_{b_{on}}$ | RB Binding Coupling | | X | 3.05 | 1.68 | (1.4, 7.32) | 0.9998 | 5,280 | 6.34 | 6.48 |
| $BR_{b_{on}}$ | BR Binding Coupling | | X | 0.13 | 1.8 | (0.05, 0.23) | 1.0001 | 5,170 | 5.51 | 6.79 |
| $R_{\gamma}$ | Rotation Current Coupling | | | 1,090 | 2.97 | (222, 7,590) | 0.9999 | 5,240 | 3.01 | 3.07 |
| $\rho_{leak}$ | Current Leakeage Ratio | | | $4.37 \cdot 10^{-7}$ | 8.7 | $(9.19 \cdot 10^{-9}, 1.06 \cdot 10^{-5})$ | 1 | 5,250 | 1.52 | 1.53 |
| $\gamma$ | Unitary Channel Current | $pA$ | | 1.93 | 1.12 | (1.61, 2.35) | 0.9999 | 5,200 | 29.2 | 28.4 |
| $i_0$ | Baseline Current | $pA$ | | 5.3 | 1.21 | (3.81, 7.21) | 0.9999 | 5,130 | 16.9 | 16.9 |
| $\epsilon^2$ | White Noise | $pA^2\, s^{-1}$ | | $2.93 \cdot 10^{-4}$ | 1.11 | $(2.47 \cdot 10^{-4}, 3.52 \cdot 10^{-4})$ | 0.9999 | 4,670 | 29.9 | 29.9 |
| $\nu^2$ | Pink Noise | $pA^2$ | | 0.61 | 1.22 | (0.43, 0.8) | 1.0007 | 4,380 | 16.8 | 16.9 |
| $N_{ch}$ | Number of Channels | | | 1,140 | 1.13 | (920, 1,370) | 0.9999 | 5,240 | 27.8 | 27.5 |
| $k_{inact}$ | Inactivation Rate | $s^{-1}$ | | $3.36 \cdot 10^{-4}$ | 1.12 | $(2.95 \cdot 10^{-4}, 3.83 \cdot 10^{-4})$ | 1.0001 | 5,410 | 29.2 | 40.6 |

**Table S2**: Characterization of the posterior distributions for Scheme X parameters. [a] Parameter symbol. [b] Full parameter name. [c] Units of measurement. [d] An "X" indicates that the de-mixing operation was applied over all samples $s$, defined as $BR(s) > RB(s) \Rightarrow RB(s) \leftrightarrow BR(s), \quad BR_{r_{\text{on}}}(s) \leftrightarrow RB_{r_{\text{on}}}(s), \quad BR_{b_{\text{on}}}(s) \leftrightarrow RB_{b_{\text{on}}}(s)$ where $\leftrightarrow$ indicates a swap operation. [e] Posterior median. [f] Geometric standard deviation (GSD), computed as $10^{\text{sd}}$, where sd is the standard deviation of the $\log_{10}$-transformed posterior; a value of 10 indicates dispersion by a factor of 10. [h] Equal-tail 90% credible interval (5%–95% quantiles). [i] Potential scale reduction factor, $\hat{R}$, used to assess convergence. [j] Effective sample size (ESS). [k] Posterior Concentration Factor based on standard deviations, defined as $PCF_{sd} = \frac{\text{sd}^{\text{prior}}}{\text{sd}^{\text{post}}}$. [l] Posterior Concentration Factor based on credible intervals, defined as $PCF_{CI} = \frac{\log(\text{prior}_{0.95}) - \log(\text{prior}_{0.05})}{\log(\text{post}_{0.95}) - \log(\text{post}_{0.05})}$. For each sample $s$, both PCF metrics quantify the increase in precision from the prior to the posterior; values greater than 1 indicate that the posterior distribution is more concentrated, reflecting reduced uncertainty upon incorporating the data.

| Parameter[a] | Full Name[b] | Units[c] | DM[c] | Median[c] | gsd [g] | $90\%CI$[e] | $\hat{R}$ [i] | ESS [j] |
|---|---|---|---|---|---|---|---|---|
| $b_{on}$ | Binding on | $\mu$M$^{-1}$ $s^{-1}$ | | 10.7 | 25.6 | (0.05, 2,010) | 1 | 5,290 |
| $b_{off}$ | Binding off | $s^{-1}$ | | 103 | 26.9 | (0.45, $2.34 \cdot 10^4$) | 1.001 | 5,270 |
| $r_{on}$ | Rotation on | $s^{-1}$ | | 95.2 | 25.6 | (0.42, $2.00 \cdot 10^4$) | 1.0001 | 5,110 |
| $r_{off}$ | Rotation off | $s^{-1}$ | | 106 | 24.2 | (0.54, $1.81 \cdot 10^4$) | 0.9999 | 5,210 |
| $RB$ | RB Equilibrium Coupling | | | 110 | 25.1 | (0.59, $2.26 \cdot 10^4$) | 1.0002 | 5,200 |
| $BR$ | BR Equilibrium Coupling | | | 9.36 | 25.9 | (0.05, 2,040) | 1.0001 | 5,090 |
| $RB_{r_{on}}$ | RB Rotation Coupling | | | 0.97 | 26.8 | ($5.03 \cdot 10^{-3}$, 241) | 0.9999 | 5,310 |
| $BR_{r_{on}}$ | BR Rotation Coupling | | | 1.01 | 27.5 | ($4.76 \cdot 10^{-3}$, 273) | 0.9999 | 5,010 |
| $RB_{b_{on}}$ | RB Binding Coupling | | | 1.05 | 26.2 | ($4.83 \cdot 10^{-3}$, 214) | 0.9999 | 5,270 |
| $BR_{b_{on}}$ | BR Binding Coupling | | | 0.98 | 25.9 | ($4.58 \cdot 10^{-3}$, 192) | 1 | 4,930 |
| $RB$ | RB Equilibrium Coupling | | X | 234 | 16.2 | (3.23, $2.90 \cdot 10^4$) | 1.0003 | 5,200 |
| $BR$ | BR Equilibrium Coupling | | X | 4.22 | 16 | (0.04, 346) | 1.0003 | 5,100 |
| $RB_{r_{on}}$ | RB Rotation Coupling | | X | 0.96 | 26.6 | ($4.94 \cdot 10^{-3}$, 222) | 1.0003 | 5,100 |
| $BR_{r_{on}}$ | BR Rotation Coupling | | X | 1.02 | 27.6 | ($4.83 \cdot 10^{-3}$, 292) | 0.9999 | 4,660 |
| $RB_{b_{on}}$ | RB Binding Coupling | | X | 1.06 | 26.5 | ($5.00 \cdot 10^{-3}$, 229) | 1.0002 | 5,120 |
| $BR_{b_{on}}$ | BR Binding Coupling | | X | 0.97 | 25.6 | ($4.50 \cdot 10^{-3}$, 186) | 1 | 4,590 |
| $R_{\gamma}$ | Rotation Current Coupling | | | 94.3 | 26.3 | (0.43, $2.22 \cdot 10^4$) | 1.0002 | 5,120 |
| $\rho_{leak}$ | Current Leakeage Ratio | | | $9.69 \cdot 10^{-7}$ | 26.9 | ($5.12 \cdot 10^{-9}$, $2.55 \cdot 10^{-4}$) | 1.0001 | 5,080 |
| $\gamma$ | Unitary Channel Current | $pA$ | | 1.1 | 26.1 | ($5.63 \cdot 10^{-3}$, 243) | 1 | 5,160 |
| $i_0$ | Baseline Current | $pA$ | | 0.98 | 25.7 | ($4.42 \cdot 10^{-3}$, 216) | 0.9999 | 4,970 |
| $\epsilon^2$ | White Noise | $pA^2\, s^{-1}$ | | $9.69 \cdot 10^{-4}$ | 24.8 | ($5.09 \cdot 10^{-6}$, 0.19) | 1.0002 | 4,990 |
| $\nu^2$ | Pink Noise | $pA^2$ | | 5.43 | 26.7 | (0.03, 1,130) | 0.9998 | 5,200 |
| $N_{ch}$ | Number of Channels | | | 4,630 | 28.2 | (20.3, $1.23 \cdot 10^6$) | 1.0002 | 5,370 |
| $k_{inact}$ | Inactivation Rate | $s^{-1}$ | | $1.02 \cdot 10^{-5}$ | 25.8 | ($4.85 \cdot 10^{-8}$, $1.95 \cdot 10^{-3}$) | 1.0006 | 4,540 |

**Table S3**: Characterization of the Affine Invariant sampling of Scheme X parameters at $\beta = 0$, that is when only the prior distribution determined the sampling. [a] Parameter symbol. [b] Full parameter name. [c] Units of measurement. [d] An "X" indicates that the de-mixing operation (see Table S2 ) was applied over all samples. [e] Prior median. [f] Prior geometric standard deviation (gsd). The expected value for the common prior variance of 2 is $10^{\sqrt{2}} \approx 25.95$ [h] Prior Equal-tail 90% credible interval (5%–95% quantiles). The expected quantiles for a median of 1 are $(4.72 \cdot 10^{-3}, 212)$. [i] Potential scale reduction factor, $\hat{R}$, used to assess convergence. [j] Effective sample size (ESS).

| Parameter[a] | Full Name[b] | Units[c] | True Value[c] | Posterior Median [d] | Posterior 90% $CI$[e] | Diagnostic[e] |
|---|---|---|---|---|---|---|
| $b_{on}$ | Binding on | $\mu$M$^{-1}$ $s^{-1}$ | 6.02 | 6.27 | (5.76, 6.8) | ✓ (Within) |
| $b_{off}$ | Binding off | $s^{-1}$ | 824 | 860 | (759, 970) | ✓ (Within) |
| $r_{on}$ | Rotation on | $s^{-1}$ | 1,680 | 1,700 | (1,570, 1,830) | ✓ (Within) |
| $r_{off}$ | Rotation off | $s^{-1}$ | 157 | 157 | (135, 192) | ✓ (Within) |
| $BR$ | BR Equilibrium Coupling | | 3.49 | 4.83 | (1.99, 8.72) | ✓ (Within) |
| $BR_{r_{on}}$ | BR Rotation Coupling | | 2.26 | 2.81 | (2.15, 3.71) | ✓ (Within) |
| $BR_{b_{on}}$ | BR Binding Coupling | | 0.13 | 0.15 | (0.06, 0.26) | ✓ (Within) |
| $RB$ | RB Equilibrium Coupling | | 56.5 | 40.8 | (25.3, 86.7) | ✓ (Within) |
| $RB_{b_{on}}$ | RB Binding Coupling | | 3.07 | 2.94 | (1.53, 6.44) | ✓ (Within) |
| $RB_{r_{on}}$ | RB Rotation Coupling | | 3,380 | 2,370 | (1,590, 3,540) | ✓ (Within) |
| $R_\gamma$ | Rotation Current Coupling | | 1,120 | 1,080 | (221, 7,540) | ✓ (Within) |
| $\rho_{leak}$ | Current Leakeage Ratio | | $4.09 \cdot 10^{-7}$ | $4.36 \cdot 10^{-7}$ | $(1.01 \cdot 10^{-8}, 1.01 \cdot 10^{-5})$ | ✓ (Within) |
| $\gamma$ | Unitary Channel Current | $pA$ | 1.92 | 1.49 | (1.27, 1.76) | ↓ Under |
| $\epsilon^2$ | White Noise | $pA^2\ s^{-1}$ | $2.49 \cdot 10^{-4}$ | $2.86 \cdot 10^{-4}$ | $(2.45 \cdot 10^{-4}, 3.37 \cdot 10^{-4})$ | ✓ (Within) |
| $\nu^2$ | Pink Noise | $pA^2$ | 0.61 | 0.55 | (0.42, 0.7) | ✓ (Within) |
| $i_0$ | Baseline Current | $pA$ | 5.3 | 5.43 | (4.18, 7.36) | ✓ (Within) |
| $N_{ch}$ | Number of Channels | | 1,140 | 1,470 | (1,220, 1,740) | ↑ Over |
| $k_{inact}$ | Inactivation Rate | $s^{-1}$ | $3.36 \cdot 10^{-4}$ | $3.4 \cdot 10^{-4}$ | $(3.04 \cdot 10^{-4}, 3.80 \cdot 10^{-4})$ | ✓ (Within) |

**Table S4**: Validation of MacroIR using simulated data. To assess the robustness of our Bayesian framework, we validated the MacroIR method using simulated recordings. We generated synthetic ATP-activated currents based on known kinetic parameters for Scheme X, ensuring that the simulation conditions matched our experimental setup. We then applied MacroIR to this dataset to evaluate its ability to recover the original parameters. [a] Parameter symbol. [b] Parameter full name. [c] Units [d] Parameter value used to generate the simulated data.[e] Median of the posterior distribution of each parameter after the MacroIR algorithm over the simulated data and the de-mixing operatorion [f] Credible Interval of each parameter. [g] Diagnostic of the recovery of each parameter. Is the recovered value within the 90% Credible Interval (green) is it overestimated (red) or is it underestimated (blue).

| Region | chain | Percentage | Stand deviation |
|---|---|---|---|
| Head-Loop | C | 9.51 | 5.85 |
| Dorsal fin | C | 85.32 | 5.08 |
| Lower Body | C | 68.09 | 15.41 |
| Left flipper | C | 59.71 | 20.96 |

**Table S5**: $P2X^{closed}_{1\text{-}ATP}$ DCOD averages (See Eq.S8) for chain C Head-Loop, LF, DF, and LB.
